# A neuromechanistic model for rhythmic beat generation

**DOI:** 10.1101/397075

**Authors:** Amitabha Bose, Áine Byrne, John Rinzel

**Author notes:** All authors contributed equally to this work.

## Abstract

When listening to music, humans can easily identify and move to the beat. Numerous experimental studies have identified brain regions that may be involved with beat perception and representation. Several theoretical and algorithmic approaches have been proposed to account for this ability. Related to, but different from the issue of how we perceive a beat, is the question of how we learn to generate and hold a beat. In this paper, we introduce a neuronal framework for a beat generator that is capable of learning isochronous rhythms over a range of frequencies that are relevant to music and speech. Our approach combines ideas from error-correction and entrainment models to investigate the dynamics of how a biophysically-based neuronal network model synchronizes its period and phase to match that of an external stimulus. The model makes novel use of on-going faster gamma rhythms to form a set of discrete clocks that provide estimates, but not exact information, of how well the beat generator spike times match those of a stimulus sequence. The beat generator is endowed with plasticity allowing it to quickly learn and thereby adjust its spike times to achieve synchronization. Our model makes generalizable predictions about the existence of asymmetries in the synchronization process, as well as specific predictions about resynchronization times after changes in stimulus tempo or phase. Analysis of the model demonstrates that accurate rhythmic time keeping can be achieved over a range of frequencies relevant to music, in a manner that is robust to changes in parameters and to the presence of noise.

**Author summary:** Music is integral to human experience and is appreciated across a wide range of cultures. Although many features distinguish different musical traditions, rhythm is central to nearly all. Most humans can detect and move along to the beat through finger or foot tapping, hand clapping or other bodily movements. But many people have a hard time “keeping a beat”, or say they have “no sense of rhythm”. There appears to be a disconnect between our ability to perceive a beat versus our ability to produce a beat, as a drummer would do as part of a musical group. Producing a beat requires beat generation, the process by which we learn how to keep track of the specific time intervals between beats, as well as executing the motor movement needed to produce the sound associated with a beat. In this paper, we begin to explore neural mechanisms that may be responsible for our ability to generate and keep a beat. We develop a computational model that includes different neurons and shows how they cooperate to learn a beat and keep it, even after the stimulus is removed, across a range of frequencies relevant to music. Our dynamical systems model leads to predictions for how the brain may react when learning a beat. Our findings and techniques should be widely applicable to those interested in understanding how the brain processes time, particularly in the context of music.

## Introduction

Humans have the ability to estimate and keep track of time over a variety of timescales in a host of different contexts ranging from sub-seconds to tens of seconds or more [1,2]. On the millisecond to second time scale, for example, numerous studies have shown that humans can accurately discriminate shorter intervals from longer intervals [3,4]. On a longer timescale, we utilize a form of time estimation that can span hours, days or years [5]. Many such examples involve the brain making a calculation over a single event, so-called “interval timing” [6,7]. Humans can also track timing that involves multiple or repeated events. For example, we instinctively move to the beat of a piece of music through a form of sensorimotor synchronization, so-called beat-based timing [8–11]. Doing so involves identifying an underlying beat within a piece of music and coordinating the frequency and timing of one’s movements to match this beat.

Understanding how humans perceive a beat has been an active area of research for quite some time. Beat perception refers to our ability to extract a periodic time structure from a piece of music. It is a psychological process in which beats can be perceived at specific frequencies, even when the musical stimulus does not specifically contain that frequency [12]. In a recent study by Nozaradan *et al*. [13], brain activity was found to entrain to the beat frequency of a musical rhythm. Additionally, participants with strong neural entrainment exhibited the best performance when asked to tap to the rhythm [13]. Various parts of the brain have been identified as being active during beat perception. Grahn and Brett reported that basal ganglia and the supplementary motor area showed increased activity for beat-based tasks, and as such, postulated that these areas mediate beat perception [14]. Interestingly, fMRI studies of participants asked to lie still with no movement while listening to music revealed that the putamen, supplementary motor area, and premotor cortex are active [15]. Thus although no external movement may be occurring, various motor areas are nevertheless active when the brain is representing a passage of time. From the theoretical perspective, error-correction [16–22], entrainment [12,23,24], and Bayesian [25–27] models have been proposed to account for the ability to perceive a beat.

Many beat perception studies have involved finger tapping while listening to a piece of music or a metronome [13,28–32]. However, humans can also mentally conjure a beat in the absence of motor movement and external stimuli. These observations, in part, lead us to ask what neural mechanisms might be responsible for detecting, learning and generating a beat. We define beat generation as the brain’s process of construction and maintenance of a clock-like mechanism that can produce repeatable, essentially constant, time intervals that demarcate a beat. In this formalism, the brain may be monitoring the firing times of neurons involved in beat generation to match event times of an external source. Beyond such reactive synchronization, we suggest that beat generation can occur as a strictly internally created and self-driven phenomenon in the absence of external stimuli.

Models of beat generation should take into consideration several empirically observed features taken from human finger tapping studies in the presence of isochronous tones (evenly spaced in time). First, the model’s output should rapidly synchronize with the external tone sequence. Second, a model should mimic the human ability to continue tapping even after the stimulus is removed [33], a property known as synchronization-continuation. Third, a model should quickly resynchronize after a tempo change or perturbation (deviant or phase shift) to the tone sequence. Fourth, ideally a model should be capable of addressing the phenomenon of negative mean asynchrony (NMA), the reported tendency of humans to tap on average prior to tone onsets. Although the cause of, extent of, and even the existence of NMA are still in dispute new models can potentially provide insight into the phenomenon.

Two primary modeling frameworks for addressing the above mentioned properties have been proposed: entrainment models and error-correction models. Entrainment models rely on principles of dynamical systems. Mathematically, entrainment refers to an external forcing that sychronizes a set of oscillators to a specific frequency. In the context of beat perception, entrainment models posit the existence of oscillators that resonate and entrain to the underlying periodicity creating an oscillation whose spectral profile matches that of the sound sequence. These models have been used to explain various beat-related phenomena including the emergence of pulse and meter [24] and the missing pulse percept [12]. These oscillator models are typically abstract mathematical formulations and, although generic in structure, presuppose a formulation in which the system is poised near to oscillatory-destabilization of a steady state, a Hopf bifurcation [24]. Error-correction models, on the other hand, are formulated at an algorithmic level to understand how a motor movement, such as a finger tap, can be synchronized to an isochronous tone sequence [18–22]. Errors between the current tap and tone times and between the current intertap time and stimulus period are used to adjust the timing of the next finger tap. Error-correction models provide different algorithmic ways in which to make an adjustment (see [29] for a review), but typically do not propose mechanisms for how a set of neurons would estimate and correct for the error.

In this paper, we introduce a neuromechanistic framework that can be used to construct neuronal network models that are capable of learning and retaining isochronous rhythms. In its simplest form, the network consists of a single, biophysically-based, beat generator neuron (*BG*), a periodic brief stimulus and a time-interval computation mechanism based on counting cycles of a gamma oscillation. The *BG* does not directly receive input from the external stimulus and is thus not being entrained by it. Instead, the *BG* learns (within a few cycles) the frequency of the stimulus thereby allowing the *BG* to continue oscillating at this frequency, even in the absence of the stimulus. Our approach combines ideas from entrainment based, information processing based and interval timing based models. In part, it extends the heuristic two-process model [18–21] to a neural setting, by developing a neural system that learns the period and phase of the periodic stimulus, in order to bring its spikes into alignment with the stimulus tones.

A central feature of our model is the concept of a gamma counter. Gamma rhythms (30-90 Hz) are ubiquitous throughout the human nervous system [33,34]. Here we utilize roughly 40 Hz gamma oscillations to form two discrete-time clocks that count the number of gamma cycles between specific events. This idea is similar in spirit to pacemaker-accumulator models, which also use counting mechanisms [6,7,35]. In our case, one clock counts gamma cycles between successive onsets of a stimulus, sometimes called the interonset interval in behavioral studies. Another neuronal clock counts gamma cycles between successive spikes of the *BG*, the interbeat interval. A comparison is made via a putative gamma count comparator (GCC) between these two counts and this information is sent to the *BG*, consistent with recent studies of neuronal circuits that can count discrete events and compare counts with stored information [36–38]. Our *BG* neuron possesses plasticity and uses the difference in count to adjust an intrinsic parameter so that it learns the interonset interval of the stimulus. The gamma counters also provide information to the *BG* about its firing phase relative to stimulus onset times, thereby allowing for the possibility of synchronization with zero-phase difference of the *BG* spikes with the stimulus. The coupling of the stimulus and *BG* counts through the GCC is both nonlinear and non-periodic. This contrasts with coupling between stimulus and oscillator in many entrainment and information processing models where either the phase or period is directly targeted for change. Further, in such models either the coupling is periodic or the update rules are linear or vice versa [39]. Our model updates are neither periodic or linear. We note that the neuronal clocks that count cycles need not operate exclusively in the range of 40 Hz. The comparison mechanism that we describe will work for any sufficiently fast frequency oscillator.

In this paper, we will show how the *BG* model learns and holds an isochronous beat over a wide range of frequencies that includes the band of0.5 Hz to 8 Hz which is relevant for beat generation and perception. Using mathematical analysis and a continuous time clock, we explain first how the *BG* learns to period match. We then show how this extends to both period and phase for the discrete time clock counters. As will be seen, the discrete time clocks give rise to a natural bounded variability of spike times of the BG even when it is holding a beat. We will test our model using standard paradigms which parallel behavioral studies that are designed to study specific aspects of beat perception and production, such as NMA [13,40], synchronization-continuation [41] and fast resynchronization times [42,43]. The BG’s mean firing time exhibits negative mean asynchrony and we describe how this NMA can be manipulated. The model resynchronizes rapidly in response to perturbations. For an abrupt change in tempo we predict and explain with the model an asymmetry in the resynchronization time for increases versus decreases in tempo. As with tempo changes, our model predicts an asymmetry in the resynchronization time due to a phase advance versus delay of the stimulus sequence. An asymmetry also arises for a transient perturbation, a single deviant onset timing in the stimulus sequence. We explain these effects by understanding how our model incorporates linear, but discrete step, error-correction to invoke non-linear changes in frequency of the BG. In turn, we develop a set of testable predictions for human behavior that help to contrast our proposed model framework from existing ones.

## Materials and methods

The main components of our model consist of a periodic stimulus with an associated neuron *S* whose spikes mark each stimulus onset, a neuronal model for the beat generator, *BG*, and a gamma count comparator, GCC, which acts as a type of neural integrator as well as error detector. These components are linked together as shown in Fig. 1A. The output from the spiking neuron *S* and of the *BG* are sent to the GCC.

**Fig 1.**
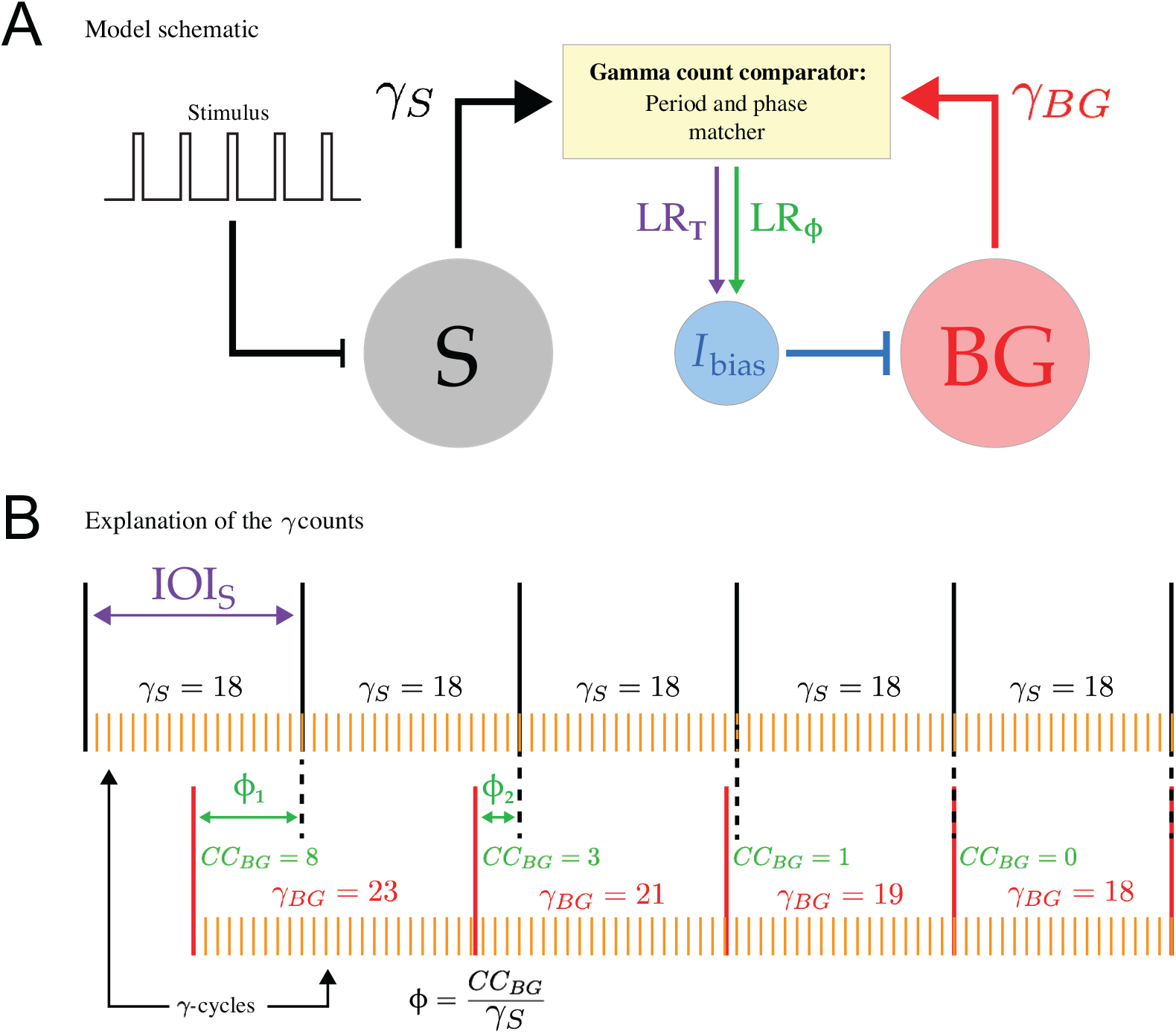
Model Schematic. **A.** The basic components of the model include a periodic stimulus, a neuron *S* whose firing demarcates the interonset interval, a beat generator neuron *BG* and a gamma-count comparator, GCC. Output from the GCC is sent via the period and phase learning rules *LR_T_* and *LR_ϕ_* to adjust *I*_bias_, which controls the frequency of the *BG*. **B.** The black vertical lines indicate periodic spike times of the *S* neuron which mark the stimulus onset. In this schematic, the interonset interval *IOI_S_* is subdivided into 18 gamma cycles (*γ_S_* = 18) as indicated. The red vertical lines indicate *BG* firing times with the gamma counts *γ_BG_* as indicated. As the *BG* spikes align to the stimulus, both *γ_BG_* and the phase *ϕ* = *CC_BG/γS_* change, until *γ_BG_* = *γ_S_* and *ϕ* = 0.

There a comparison is made which is then sent via a period learning rule, *LR_T_*, and a phase learning rule, *LR_ϕ_*, to adjust *I*_bias_ which subsequently changes the instantaneous frequency of the *BG*. The term *I*_bias_ is taken here to represent the drive to the *BG* that governs its frequency. It could be considered as a parameter internal to the *BG*, or can be more generally associated with summed synaptic input that drives the *BG*. In either case, it is a term that regulates the *BG*’s excitability.

### The beat generator and stimulus

The *BG* in our model can be described using biophysical, conductance based equations and is required to have only two specific properties. The first is that it possesses a sharp (voltage) threshold. A spike of the *BG* occurs when the voltage increases through this threshold. The second requirement is that the *BG* has a slow process that governs the time between its spikes. A simple model that possesses these basic properties is the leaky integrate and fire (*LIF*) model which we first use to describe our analytic findings. Our simulation studies utilize a biophysical model motivated by models of delta waves in sleep, as we require a similar frequency range for the *BG*. To that end, we chose voltage-gated currents similar to those from an idealized model for sleep spindle rhythms of thalamo-cortical relay cells, namely for the slow wave of relay cells in burst mode [44]. The slow wave can be generated by the interplay of a *T*-type calcium current, and *I_h_* current and a leak current. Here, the *BG* has a persistent sodium *I*_NaP_, *T*-type calcium *I*_CaT_, sag *I_h_* and leak *I_L_* currents. In the text, we refer to this as the *I_NaP_* model; for a full description and the equations see Appendix. The analytic results also hold for the *I_NaP_* model, but as the analysis is more complicated, showing so is outside the scope of this paper. For either the *LIF* or *I_NaP_* models, parameters are chosen that allow for a wide range of intrinsic frequencies of the *BG* up through 8 Hz as *I*_bias_ is varied as quantified by a neuron model’s *I*_bias_ versus frequency relationship which is presented in the Results. This range of frequencies is appropriate for speech and music. The time between successive spikes of the *BG* is called the interbeat interval and is denoted by *IBI_BG_*.

The voltage in the *LIF* model evolves according to

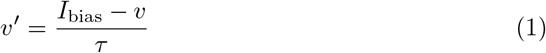

where *v* is a dimensionless variable representing voltage, *I*_bias_ is the drive to the neuron and τ is the membrane time constant. The *LIF* model has a spike and reset condition which makes it discontinuous. When the voltage reaches one at *t* = *t_s_*, it is instantaneously reset to the value 0; if 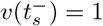, then 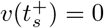. When *I*_bias_ > 1 oscillations exist. In this case, the *LIF* model is rhythmic with period given by

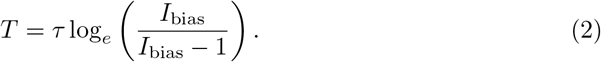

The period of *BG* given by equation (2) can be adjusted to any positive value by appropriately adjusting *I*_bias_.

For both the *LIF* and *I_Nap_* models, the specific nature of the stimulus is not modeled, only the onset is of interest here. We limit our simulations to a range between 1 and 6 Hz, which corresponds to an interstimulus interval ranging from1 s down to 166 ms. There is no theoretical or practical problem to extend the model outside of this range, as further addressed in the Discussion. We utilize a neuron *S* to faithfully transform the stimulus sequence into spikes. The interonset interval, *IOI_S_*, is then defined as the time between successive *S* spikes. The model for *S* is not important provided that it is set to be an excitable neuron that fires quickly in response to input; see the Appendix for equations.

### The gamma oscillation counters and learning rules

The gamma count comparator, GCC, in our model utilizes two generic oscillators with frequency sufficiently larger than that of both the stimulus and the *BG*. Here it is taken to lie in the gamma range at roughly 40 Hz (Fig. 1B). We choose the oscillators to be identical, though this is not a requirement of the model. To avoid integer values, both have a frequency of 36.06 Hz (period 27.73 ms); see the Appendix for details. We let *γ_BG_* be a variable that counts the number of gamma cycles between consecutive *BG* spikes and *γ_S_* be a variable that counts the number of gamma cycles between consecutive spikes of *S*. At each spike of *BG* or *S*, the appropriate counter is reset to zero. We stress that *γ_BG_* and *γ_S_* are integers, but, in general, the periods that they are estimating are not integer multiples of a gamma cycle (27.73 ms). Hence, although the stimulus period may be constant the gamma counts may vary from cycle to cycle. The difference *γ_BG_* − *γ_S_* provides an estimate of how different or aligned the frequencies of *BG* and *S* are. For example, if *S* oscillates at 5 Hz and the *BG* is initially oscillating at 3 Hz, then the GCC would count roughly 8 cycles between *S* spikes and 13 cycles between *BG* spikes. In this case, the GCC determines that the *BG* is oscillating too slowly and sends a speed up signal to the *BG*. Alternatively if the *BG* were initially oscillating at 6 Hz, then the GCC counts roughly 6 cycles and sends a slow down signal. In general, speeding up or slowing down of the *BG* is achieved by changing *I*_bias_. At each spike of the *BG*, the period learning rule adjusts *I*_bias_ by a fraction of the difference between the gamma oscillator count of cycles between *S* and *BG*. This learning rule (*LR_T_*) assigns at each *BG* spike

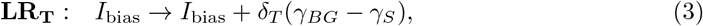

where the parameter *δ_T_* is independent of period. This simple rule is enough to align the frequencies of *S* and *BG*, the details of which will be explored through the derivation and analysis of a one-dimensional map. However, this frequency matching rule does not provide the beat generator with any information about its firing phase relative to stimulus onset.

To align the phase of the *BG* to the stimulus onset, we formulate a second learning rule. We define the current count of the *BG*, *CC_BG_*, as the number of gamma cycles from the last *BG* spike to the current *S* spike; see Fig. 1B. Then at each *S* spike define *ϕ* = *CC_BG_/γ_S_* to be the phase of *BG* firing. We use the phase to determine if the *BG* fires “before” or “after” *S* at each cycle. In a rhythmically active network, the concept of whether *BG* fired before *S* is somewhat ambiguous. We define the *BG* to be “before” the stimulus if it fires in the second half of the stimulus period *ϕ* ∈ (0,0.5). In this case we say that the *BG* is too fast and needs to slow down. Conversely, if *ϕ* ∈ (0.5,1), the *BG* is said to fire “after” *S* and needs to be sped up. At each *S* spike, we update *I*_bias_ with the second part of the learning rule (*LR_ϕ_*)

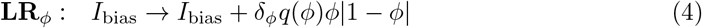

where *δ_ϕ_* is independent of period and phase and *q*(*ϕ*) = *sgn*(*ϕ* − 0.5), with *q*(0.5) < 0. Thus if *ϕ* = 0 (or 1), there is no change to *I*_bias_. But if the *BG* fires before *S* (*ϕ* ∈ (0,0.5)), then *q*(*ϕ*) < 0 and *I*_bias_ is decreased to slow down the *BG*. The opposite occurs if the *BG* fires after *S*. The absolute value keeps the last term positive as *ϕ* can become larger than 1. For example, during transitions from high to low frequency, *CC_BG_* can exceed *γ_S_*. The quadratic nature of *LR_ϕ_* is chosen so that the maximum change occurs for phases near 0.5. With this two-part learning rule, the *BG* learns both the period and phase of *S*. Both parts of the rule are implemented concurrently so that the process of period and phase alignment occurs simultaneously. The two rules *LR_T_* and *LR_ϕ_* target the value *I*_bias_. In the Results section, we describe how these changes to *I*_bias_, in turn, affect the frequency of the *BG* which then affects the period and phase of oscillations.

### Synchronizing to the beat, stationary behavior and natural variability of spike times

Given the discreteness of our gamma counters, the *BG* learns to fire a spike within a suitably short window of time of the stimulus onset, an interval equal to plus or minus one gamma cycle. We define this concept as *one gamma cycle accuracy*. For the earlier described choice of parameters, this amounts to ±27.73 ms from stimulus onset. We address two important and related concepts: synchronization to the beat and holding a beat. In our model, synchronization to the beat refers to the process by which the *BG* brings its spike times within one gamma cycle accuracy of a specific stimulus frequency. Holding a beat refers to the ability of the *BG* to maintain synchronized firing at a specific frequency over a specified stretch of time. We will say that *BG* has synchronized to the stimulus if three consecutive *BG* spikes each fall within one gamma cycle accuracy in time of a stimulus onset. The *BG* is said to be holding a beat for as long as it continues to remain synchronized with the stimulus onset.

In the presence of an isochronous stimulus, the *BG* displays what we shall call stationary behavior. This refers to the pattern of spike times of the *BG* in response to a fixed frequency stimulus. Despite there being no source of noise in our model, the discrete nature of the gamma count comparator allows the *BG*’s spike times during stationary behavior to naturally display variability. Thus, during stationary behavior, while the *BG*’s spike times typically land within one gamma cycle accuracy of stimulus onset they can also fall outside this window. The variability of the *BG*’s spikes arises because the gamma counters and learning rules adjust *I*_bias_ in discrete steps whenever *γ_S_* = *γ_BG_* or *ϕ* = 0. What this means is that during stationary behavior, the *BG* does not converge to a limit cycle oscillation (periodic orbit). The variables that govern the dynamics of the *BG* do not periodically return to the same values, but instead can vary by small amounts from cycle-to-cycle. In practice, these small differences affect the exact spike times of the *BG*, creating the variability. Furthermore, as the gamma counts are not exact representations of the period, they may be equal even when *IOI_S_* and *IBI_BG_* are unequal. This amounts to additional variability in the *BG*’s spike times relative to the spike times of *S*.

We will determine the time that it takes for the *BG* to resynchronize its spikes to stimulus onset after a change to the stimulus. Resynchronization is declared similarly to synchronization in that the *BG* is required to fire three consecutive spikes each of which must lie within one gamma cycle accuracy of a stimulus onset. The resynchronization time is then taken as the time of the first synchronized spike. In all studies, we begin with the *BG* displaying stationary behavior at a specific frequency. Because of the variability present in stationary behavior, the resynchronization times will depend on the initial conditions at the moment that the change to the stimulus profile is enacted. We will compute mean resynchronization times and standard deviations over 50 realizations, each of which differs by a small change in the initial condition of the *BG* at the time that the stimulus profile is changed. All simulations are carried out in MATLAB with a standard Euler solver (Euler-Maruyama when noise is introduced).

## Results

We provide a short outline of the results that follow. We start with a demonstration of how the *BG* learns to synchronize to an isochronous stimulus sequence. We then describe how the *BG* learns a period by first utilizing a continuous time version of the gamma counters to derive a one-dimensional map. The discrete gamma counters are then used to describe period and phase matching. Next, we present the basic behaviors of the *BG* describing its response under both stationary (isochronous stimuli) and transient (tempo changes, phase shifts and deviants) conditions. The section concludes with a brief description of the effects of parameter changes and noise.

### *I*_bias_ determines the *BG*’s frequency

An oscillatory neuronal model spikes with a period that is quantifiable by its frequency versus *I*_bias_ relation (*f-I*). This relationship is obtained from the reciprocal of (2) for the *LIF* model and computed numerically for the *I_NaP_* models (Fig. 2A). The blue (red) curve depicts the *f-I* curve for the *LIF* (*I_NaP_*) model. In the *LIF* model, the interspike interval is governed by the difference between *I*_bias_ and the spiking threshold, as well as the parameter *τ*. In the *I_NaP_* model, the interspike interval is determined by an interplay of the various non-linear currents (Fig. 2B). In particular, the *I_CaT_* and *I_L_* currents provide basic excitability to the model, the *I_NaP_* current allows for spikes once a voltage threshold is crossed and the *I_h_* current provides a slow depolarization of the membrane allowing the neuron’s voltage to gradually reach spiking threshold. Thus the primary determinant of the interspike interval is the time constant of the *I_h_* current. An important point regarding the *f-I* relations is that they are both strictly increasing. Hence, there is exactly one value of *I*_bias_ that yields a specific frequency. The learning rules we use make discrete changes to *I*_bias_. Thus, there is little chance of adjusting *I*_bias_ to the exactly correct value. Instead, the learning rules adjust *I*_bias_ so that it stays within a small window of the correct one. The frequency relations increase steeply from frequency equal to zero. Therefore, at low frequencies, larger changes in frequency can result from small changes in *I*_bias_. The same is also true for the *I_NaP_* model at frequencies in the 3 to 8 Hz range. It is important to note that any implementation of a *BG* model with a monotone increasing *f-I* relation will produce the qualitative results described below. However, the quantitative details will certainly depend on the slope and non-linearities of these relations that are produced by different ionic currents and parameters. For example, changes to *I*_bias_ in the *LIF* at frequencies above 1 Hz lead to linear changes in *BG* frequency. Alternatively, for the *I_NaP_* model a change of *I*_bias_ from say 5 to 10 produces a smaller change in *BG* frequency than changes from 10 to 15.

**Fig 2.**
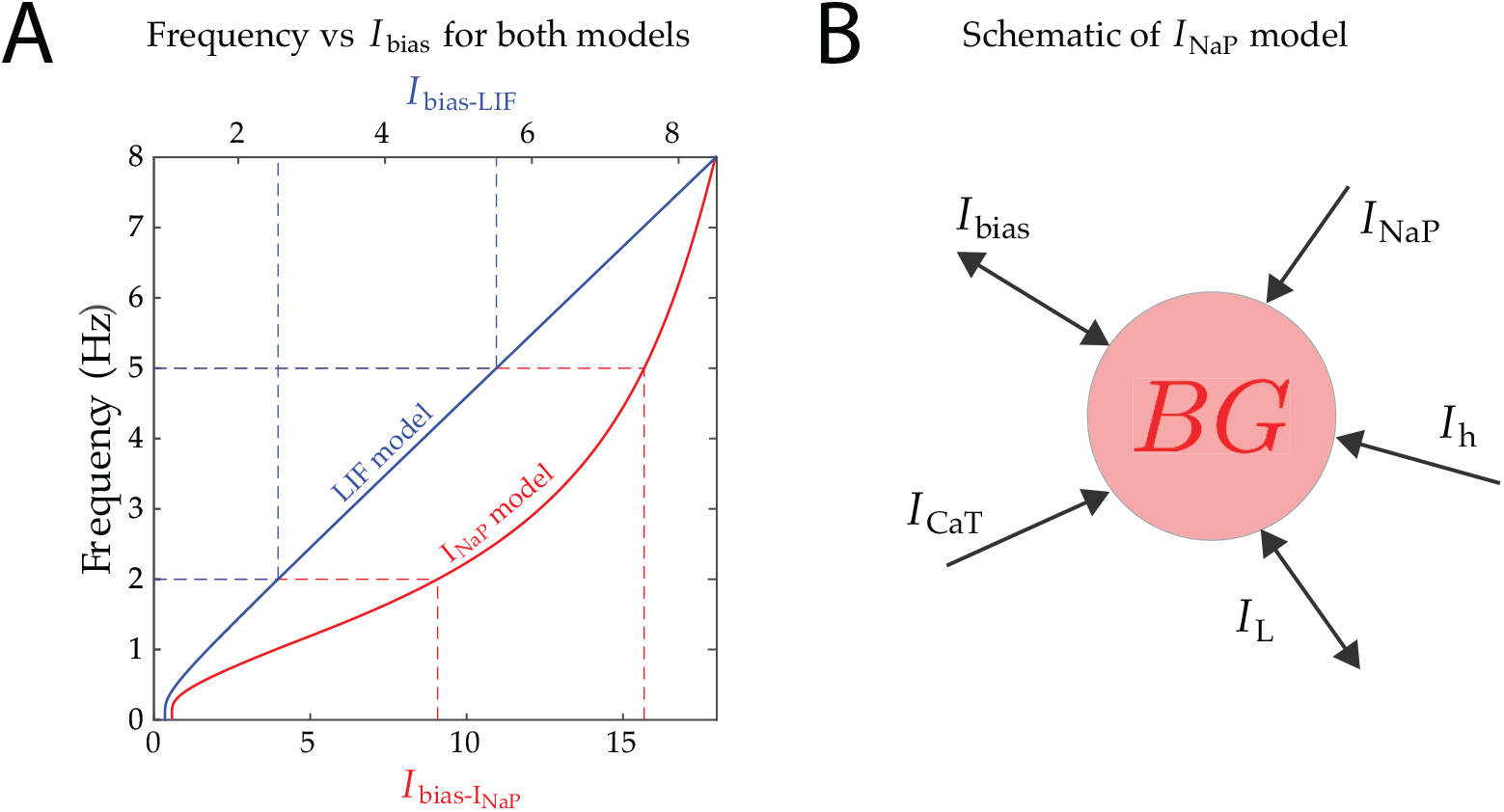
Biophysical models. **A.** Frequency vs. *I*_bias_ curves for *LIF* (blue) and *I_NaP_* (red) models. Observe the different scales for the upper and lower horizontal axes. Dashed lines show the corresponding *I*_bias_ values for two different frequencies. Both curves are increasing, starting from zero frequency, which yields qualitatively similar behavior from either model. While the *LIF* relation is almost linear for frequencies larger than 1 Hz, the *I_NaP_* relation is more power-like, with a steeper gradient from 3 to 8 Hz, this yields differences in the quantitative behavior of the models. **B.** Schematic of *I_NaP_* neuron with the different ionic currents that contribute to its excitability and spiking behavior.

The *BG* learns to oscillate at a frequency by adjusting its bias current through the set of plasticity rules *LR_T_* and *LR_ϕ_* (Fig. 3). The *BG* is initially set to oscillate at 2 Hz with *I*_bias_ = 9.06 Hz. At *t* = 0 ms, we adjust the stimulus frequency to 4.65 Hz and activate the period learning rule *LR_T_* (Fig. 3A). Notice how the cycle period of the *BG* increases on a cycle-by-cycle basis until it matches the stimulus period. This results from the value of *I*_bias_ iteratively increasing over the transition, based on the difference *γ_BG_* − *γ_S_*. The first change to *I*_bias_ does not occur until *t* = 500 ms, which is the time the *BG* naturally would fire when oscillating at 2 Hz, since *LR_T_* updates (purple curve in the bottom panel of Fig. 3A) are only made at spikes of the *BG*. At around *t* = 2.25 s, the value of *I*_bias_ falls within one gamma cycle accuracy as depicted by the blue band but continues to adjust. Note that *I*_bias_ does not settle down to a constant value. Instead it changes by ±*δ_T_* whenever |*γ_BG_* − *γ_S_* | = 1. Additionally, since *LR_T_* contains no phase information, the spikes of the *BG* are not synchronized in phase with those of *S*. At *t* = 4.2 s (20 cycles of the stimulus), the stimulus is completely removed, and the *BG* continues to oscillate at roughly 4.65 Hz. This shows that the *BG* has learned the new frequency and does not require periodic input to make it spike. There are still adjustments to *I*_bias_ because the *BG* continues to compare *γ_BG_* with the last stored value of *γ_S_*. This example demonstrates how the *BG* oscillates at a learned frequency rather than through entrainment to external input.

**Fig 3.**
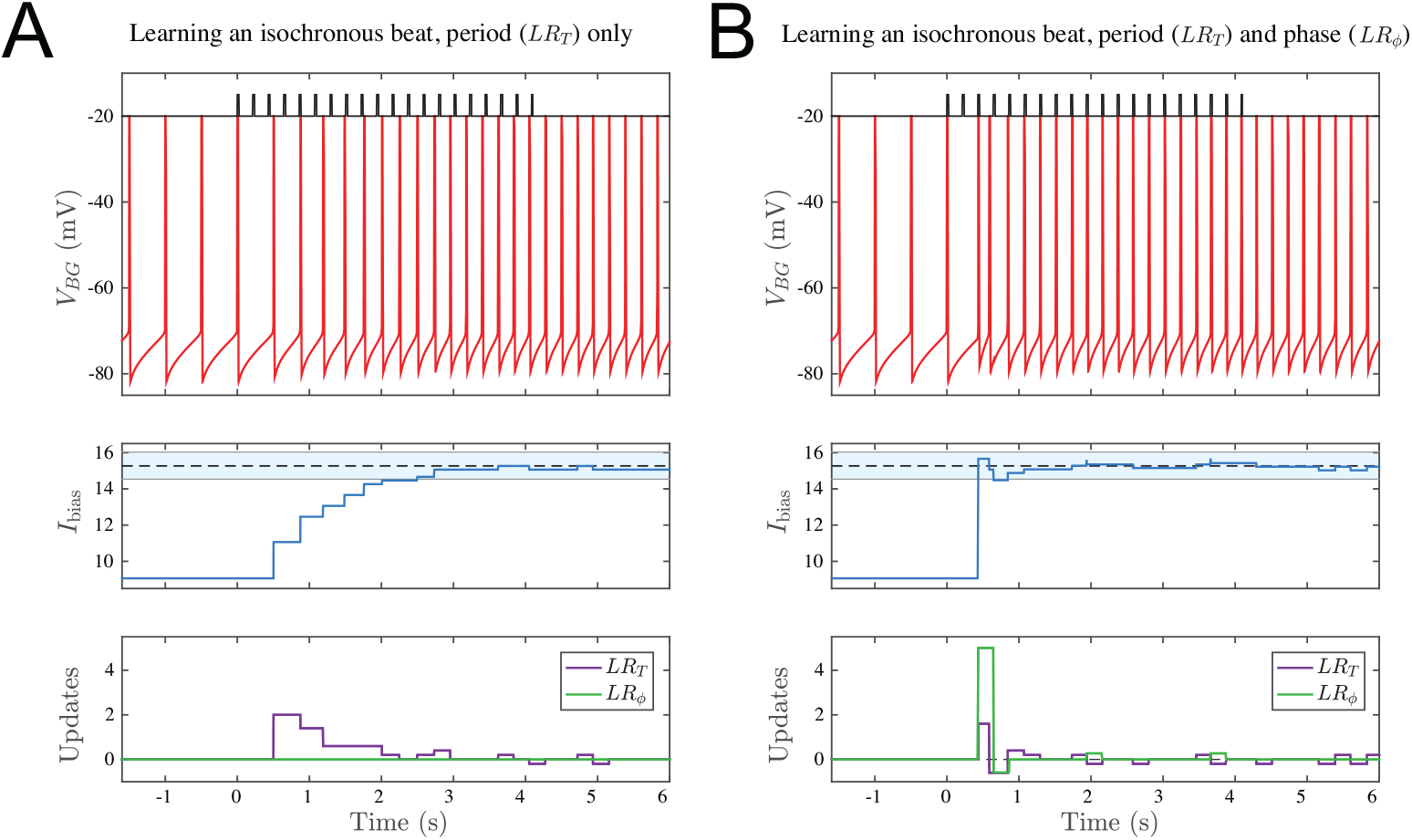
Effect of learning rules. **A.** Period Matching with *LR_T_*. The voltage time course (top) of the *BG* is shown under three different conditions; i) Endogenously oscillating at 2 Hz with no stimulus and no learning rule from *t* = −1.5 to 0 s; ii) Learning the 4.65 Hz isochronous period in the presence of the stimulus from *t* = 0 to 4.2 s; iii) Performing synchronization continuation once the stimulus is removed at *t* = 4.2 s. *I*_bias_ changes on a cycle-by-cycle basis (middle panel). Around *t* = 2.25 s, *I*_bias_ enters the shaded blue regime (14.54,16.03) that represents the one gamma cycle accuracy range from the 15.27 value that produces exactly 4.65 Hz. After *t* = 2.75 s, *I*_bias_ lies very close to the correct value. Updates to *I*_bias_ rely solely on *LR_T_*, the period learning rule (lower panel). **B.** Period and phase matching with both *LR_T_* and *LR_ϕ_*. Time courses are the counterparts of those in Panel A. Now the convergence to the correct frequency and phase occurs very quickly by about *t* = 1.2 s. Note the early large change in *I*_bias_ due to *LR_ϕ_*. In both panels after *t* = 4 s, there are still changes to *I*_bias_ as there are minor adjustments to *I*_bias_ though the frequency of the *BG* stays within the equivalent of one gamma cycle accuracy of 4.65 Hz (*IBI_BG_* = 215 ± 27 ms). Note that once the stimulus is switched off, only *LR_T_* continues to update *I*_bias_. Here, and in Figs. 5–8, we used the *I_NaP_* model and set *δ_T_* = 0.2, *δ_ϕ_* = 2.5.

When both learning rules operate together, the *BG* learns both the correct period and phase. Starting with the same initial conditions as in Fig. 3A, now at *t* = 0 ms both *LR_ϕ_* and *LR_T_* are turned on (Fig. 3B). Note the very rapid synchronization of the *BG*’s spikes with the stimulus onset times. The middle panel shows how *I*_bias_ grows much more quickly when both rules are applied. The first update is due to the phase learning rule at the third stimulus spike, at *t* = 433.5 ms, which is earlier than in the previous example. This causes enough of an increase in *I*_bias_ for the *BG* to immediately fire, which causes an update due the period learning rule. The lower panel shows how the two rules *LR_T_* and *LR_ϕ_* contribute to the change in *I*_bias_. Note that the learning is not sequential with period learning preceding phase learning or vice versa. Rather, period and phase learning occurs concurrently. Below we shall describe in more detail each of these learning rules and their role in synchronizing the *BG* with the stimulus.

### *LR_T_*: The learning rule for period matching

The dynamics of how the period learning rule *LR_T_* matches the interbeat interval of the *BG*, *IBI_BG_*, with the interonset interval of *S*, *IOI_S_*, can be explained in terms of an event-based map. Each spike of the *BG* is treated as an event and we define a map that updates the bias current *I*_bias_ on a cycle-by-cycle basis. To derive the map, we first use exact time differences, in effect, equivalent to a continuous time-keeping mechanism. This will allow the map to possess a parameter-dependent asymptotically stable fixed point. For simplicity of presentation, we use the *LIF* model to derive the specifics of the map. We will then discuss how those findings inform simulations of the *I_NaP_* model for the discrete gamma count case.

Assume that the stimulus sequence occurs with a fixed period *T*\* corresponding to a specified *IOI_S_*, and that the *BG* is initially oscillating with an interbeat interval of *T*_0_. This *IBI_BG_* corresponds to a specific value *I*_0_ of *I*_bias_ given by solving (2). *I*_bias_ is then updated to *I*_1_ by comparing *T*_0_ to *T*\*. In turn, this produces a new cycle period *T*_1_ and so on. In general, the continuous time version of *LR_T_* updates *I*_bias_ at each firing of the *BG* as follows:

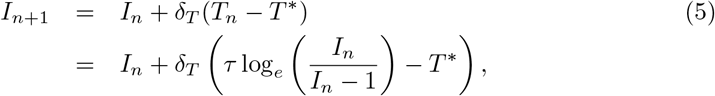

where the second line is obtained by substituting equation (2) evaluated at *I_n_* for *T_n_*. Error-correction models also take the form of an iteration scheme, but typically target the next cycle period for adjustment, i.e. *T_n_* and *T*_*n*+1_ would replace *I_n_* and *I*_*n*+1_, respectively, in the first equation of (5). In contrast, the adjustment in our model is made to the biophysical parameter *I*_bias_ (*I_n_*) which then has a subsequent effect on the cycle period (*T_n_*).

Equation (5) defines a one-dimensional map which can be expressed as *I*_*n*+1_ = *f* (*I_n_*), where *f* (*I*) denotes the right-hand side. A fixed point of the map satisfies *I* * = *f* (*I*\*) whose stability can be determined by checking the condition |*f*’(*I*\*)| < 1. A fixed point of the map corresponds to a case where the *IBI_BG_* of the *BG* is equal to the *IOI_S_* of *S*. Stability of the fixed point implies that the learning rule is convergent. Note that for any *T*\*, there is a unique fixed point of the map which satisfies *I* * = 1/(1 − exp(−*T* */*τ*)). This means that any stimulus period can in practice be learned by the *BG*, provided that the fixed point is stable. A simple calculation shows that |*f*’(*I*\*)| < 1 provided 0 < *δ_T_* < 2*I*\*(*I*\* − 1)/*τ*. For fixed *δ_T_*, as the stimulus frequency gets smaller, *I*\* converges to 1, and as a result the term 2*I*\*(*I*\* − 1)/*τ* goes to zero. This expression provides the insight that convergence for lower stimulus frequencies requires taking smaller increments in the learning rule. This finding carries over to any *f-I* relation that is steeply sloped at low frequencies.

Parameter dependence and the ensuing dynamics of the map are readily illustrated graphically (Fig. 4). The one-dimensional map has a vertical asymptote at *I* = 1, a local minima at 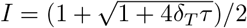 and a slant asymptote of *I* − *δ_T_*/*T*\*. The graph intersects the diagonal at exactly one point, and the slope of the intersection determines the stability as calculated above. For increasing stimulus frequency, with *δ_T_* and *τ* fixed, the map’s graph shifts upward (Fig 4A) and the fixed point moves to larger values of *I*_bias_. Note, for low stimulus frequency (here, 1 Hz) the fixed point is unstable. The update parameter *δ_T_* does not change the value of the fixed point *I*\*, but affects the stability (Fig. 4B). As *δ_T_* increases, the slope at the intersection decreases through 0, then through -1, at which point stability is lost. If the stimulus frequency changes (eg, 2 Hz to 5 Hz), *I*_bias_ changes dynamically as the *BG* learns the new rhythm. The learning trajectory corresponds to the cobweb diagram on the map (Fig. 4C, black dashed lines and arrows). Each adjustment of *I*_bias_ occurs at a spike of *BG* and allows it to speed up for the next cycle. In this example, it takes only a few cycles for the *BG* to learn the new rhythm. The transition from 5 to 2 Hz (Fig. 4C, red dashed lines and arrows) demonstrates the asymmetry in convergence for similar sized changes of opposite directions. Here the convergence for the decrease in frequency occurs over fewer cycles because the value of *δ_T_* chosen yields a fixed point at 2 Hz with near zero slope. The smaller in magnitude the slope, the less the number of cycles needed to converge. This result suggests that certain preferred frequencies can exist for specific choices of parameters.

**Fig 4.**
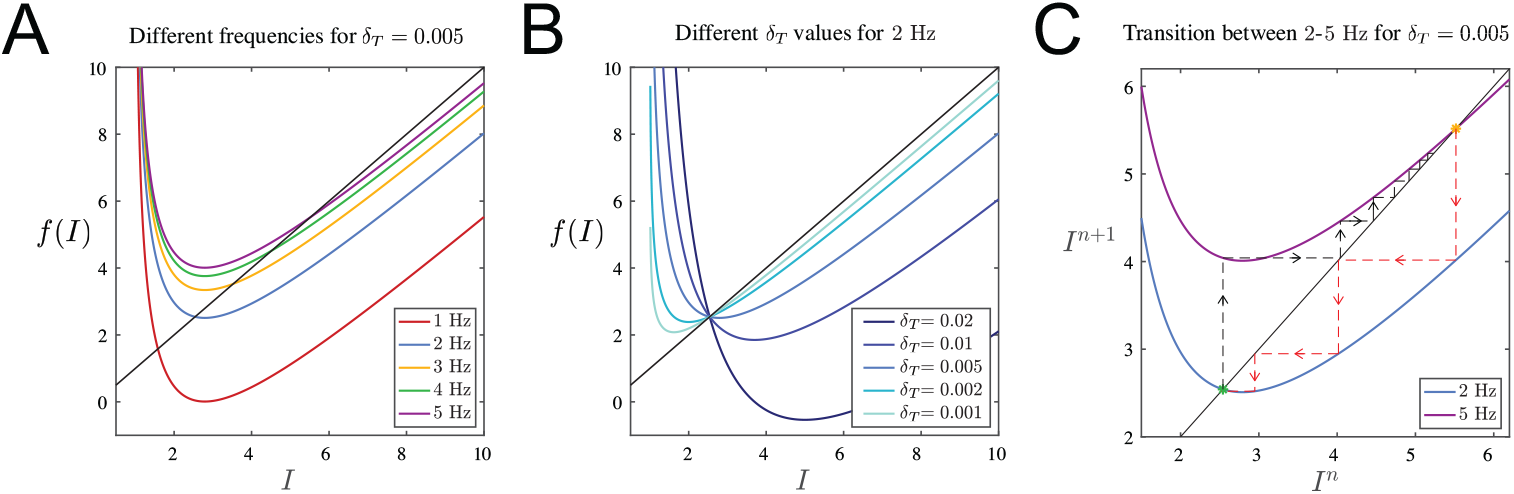
One-dimensional map for period matching illustrated using the *LIF* model. **A.** Plots of the map for different frequency stimulus are shown. Each curve crosses the diagonal at exactly one point, corresponding to a fixed point of the map. The slope at this intersection determines stability. As frequency increases, the fixed point moves up and gains stability. **B.** For the case of 2 Hz, as *δ_T_* increases, stability is lost. Similar results hold for any stimulus frequency. **C.** A cobweb diagram of the convergence of a trajectory is shown. Initially, the black trajectory starts at a value corresponding to a 2 Hz oscillation. Then *I*_bias_ is shifted to a value corresponding to a 5 Hz oscillation. The trajectory cobwebs over a number of cycles until it converges to the new stimulus frequency. The opposite transition from 5 Hz to 2 Hz is shown in red and occurs over less cycles.

In contrast to the idealized continuous-time learning rule, the gamma count-based case does not lead to updates that converge to zero. An interesting illustration is seen after an *IOI_S_* has been learned and the stimulus is turned off. Small updating persists (e.g., Fig. 3A, bottom panel). Just after the turn-off, the *IBI_BG_* is less than the last stored *IOI_S_* (*γ_BG_* < *γ_S_*). So the *BG* is too fast, and at the next *BG* spike the period rule *LR_T_* activates and decreases *I*_bias_ by *δ_T_* producing a new, longer *IBI_BG_*. Not immediately, but after a while (just after *t* = 5*s*) a difference in gamma counts again arises. This time *LR_T_* increases *I*_bias_, shortening *IBI_BG_* and so on. These changes are all due to *LR_T_* as the phase learning rule *LR_ϕ_* can never be invoked since the stimulus is no longer present.

### *LR_ϕ_*: The learning rule for phase matching and synchronization with the stimulus onset

The phase learning rule *LR_ϕ_* considers the current *BG* gamma count, *CC_BG_*, at each firing of *S*. As a result, the *BG* has information about its phase at each stimulus onset. We use a learning rule function *ϕ*|1 − *ϕ*| that has maximal effect at *ϕ* = 0.5 and no effect at *ϕ* = 0 and 1. This is similar to a logistic function that attracts dynamics towards *ϕ* =1; see also [45] for a similar mathematical rule used in a different biological context. In our case *ϕ* = 0 is equivalent *ϕ* = 1, so our learning rule *LR_ϕ_* utilizes a sign changing function *q*(*ϕ*) = *sgn*(*ϕ* − 0.5), *q*(0.5) = −1 to stabilize *ϕ* = 0 as well. This will allow convergence via either phase increase or decrease towards synchrony. At each *S* spike-time, the *BG* is sped up (if *ϕ* ∈ (0.5,1)) or slowed down (if *ϕ* ∈ (0, 0.5)) by adjusting *I*_bias_ until the phase reaches a neighborhood of 0 or 1. This, in conjunction with *LR_T_* which equalizes the *IOI_S_* and *IBI_BG_*, brings about synchronization. Note that when the *BG* fires within one gamma cycle accuracy of *S*, *ϕ* = 0 or 1. In that case, there is no update to *I*_bias_. Thus as with *LR_T_*, because of the discreteness of the learning rule updates, the value of *I*_bias_ is brought into close proximity of the value of *I*_bias_ that produces a specified rhythm but need not become exact. The rapid synchronization results shown earlier in Fig. 3 hold for a large range of stimulus frequencies. Under certain conditions, it is possible to derive a two dimensional map that tracks how *I*_bias_ and *ϕ* change on a cycle-by-cycle basis. Though it is outside the scope of this paper, an analysis of the map shows that stable period and phase matching can be achieved for each *IOI_S_*, if *δ_T_* and *δ_ϕ_* are not too large. In practice these two parameters should be chosen so that the changes due to *LR_T_*, *δ_T_* (*γ_BG_* − *γ_S_*), and *LR_ϕ_*, *δ_ϕ_ϕ*|1 − *ϕ*|, are of the same order of magnitude.

If the *IBI_BG_* is at least one gamma cycle longer than the *IOI_S_*, then *ϕ* > 1. We could have restricted the phase to be less than one by periodically extended *LR*_*ϕ*_ beyond the unit interval, but this would allow for stable fixed points at *ϕ* = 2, 3,4,etc.. Instead, the learning rule utilizes an absolute value around the term 1 − *ϕ* to keep it positive. This introduces an asymmetry in the resets of *I*_bias_. For example if *ϕ* = 0.1 then *ϕ*(1 − *ϕ*) = 0.09 but if *ϕ* = 1.1, then *ϕ*|1 − *ϕ*| = 0.11. Thus when the *BG* fires after the *S* spike after a long *IBI_BG_* (e.g. *ϕ* = 1.1) then *I*_bias_ is increased more than it is decreased if it fires before the *S* spike (e.g. *ϕ* = 0.1). As a result, the learning rule favors the *BG* firing *before* the stimulus onset, as if in anticipation. This is more pronounced at lower frequencies where the slope of the *f-I* curve is much steeper than linear. This issue is explored in more detail in the following sections.

The phase learning rule *LR_ϕ_* adjusts *I*_bias_ as opposed to directly affecting the phase of the *BG*, for example, via a perturbation and reset due to the phase response curve (PRC). Using a PRC to adjust phase would lead to a situation of entrainment rather than learning. Indeed, with a PRC, bringing the value of *I*_bias_ to within one gamma cycle accuracy to achieve a specific frequency is rarely reached. Thus if the stimulus were to be removed, a *BG* with a PRC-based phase rule would fail to continue spiking at the correct target frequency, i.e. it would fail in a synchronization-continuation task.

### Stationary behavior and the dynamics of holding a beat

To hold a beat, the *BG* must fire spikes within a time window of one gamma cycle accuracy of stimulus onset. As discussed earlier, the discreteness of the gamma counters and comparator causes the *BG* spike times to naturally display variability. Thus the *BG* must at each firing compare its period and phase relative to stimulus onset times and make necessary corrections. Holding a beat is an example of stationary behavior of the *BG* in response to a constant frequency stimulus (Fig. 5). In this typical example, here shown at 2 Hz, each spike of the *BG* is aligned to the closest spike of *S* and then a timing error equal to the *BG* spike time minus *S* spike time is computed. The value of *I*_bias_ hovers around the dashed black line *I*_bias_ = 9.06 which is the value that produces exactly a 2 Hz oscillation (Fig. 5A, upper). The spike times of the *BG* jitter around those of *S*, and thus, the timing error is poised around zero (Fig. 5A, lower). While holding a beat, these differences fall within a single gamma cycle time window (dashed gray lines at ±27.73 ms). During some time windows (pink shaded region in Fig. 5B) no updating of *I*_bias_ occurs (lower time course), but the timing differences progressively decrease and become more negative. The *BG* spike times are drifting relative to onset times because the *IBI_BG_* is slightly smaller than *IOI_S_*, but close enough that *γ_BG_* = *γ_S_*. The drift represents the fact that during this interval, the *BG* is not entrained to the stimulus. The slope of the timing error in this interval is shallower when /bias is closer to 9.06, as shown for example in the time window between *t* = 72 and 76 s. During some intervals (e.g. shaded blue region), the update rules are actively trying to keep the period and phase of the *BG* aligned with *S*. Although the timing errors are not large in this case, the counts *γ_BG_* and *CC_BG_* differ from *γ_S_* thus causing *LR_T_* and *LR_ϕ_* to be invoked. During either drifting or corrective behavior, the *BG* spike times occur closely in time with the stimulus onsets. Note that drifting and corrective behaviour continue to occur after the stimulus is turned off, but only the period rule will be active (see Fig. 3). Many studies have pointed towards drifting dynamics in human tapping experiments [46–48]. Dunlap first noted this behavior, stating that the errors tend to get progressively more negative/positive, until a correction occurs and causes a change in direction [46].

**Fig 5.**
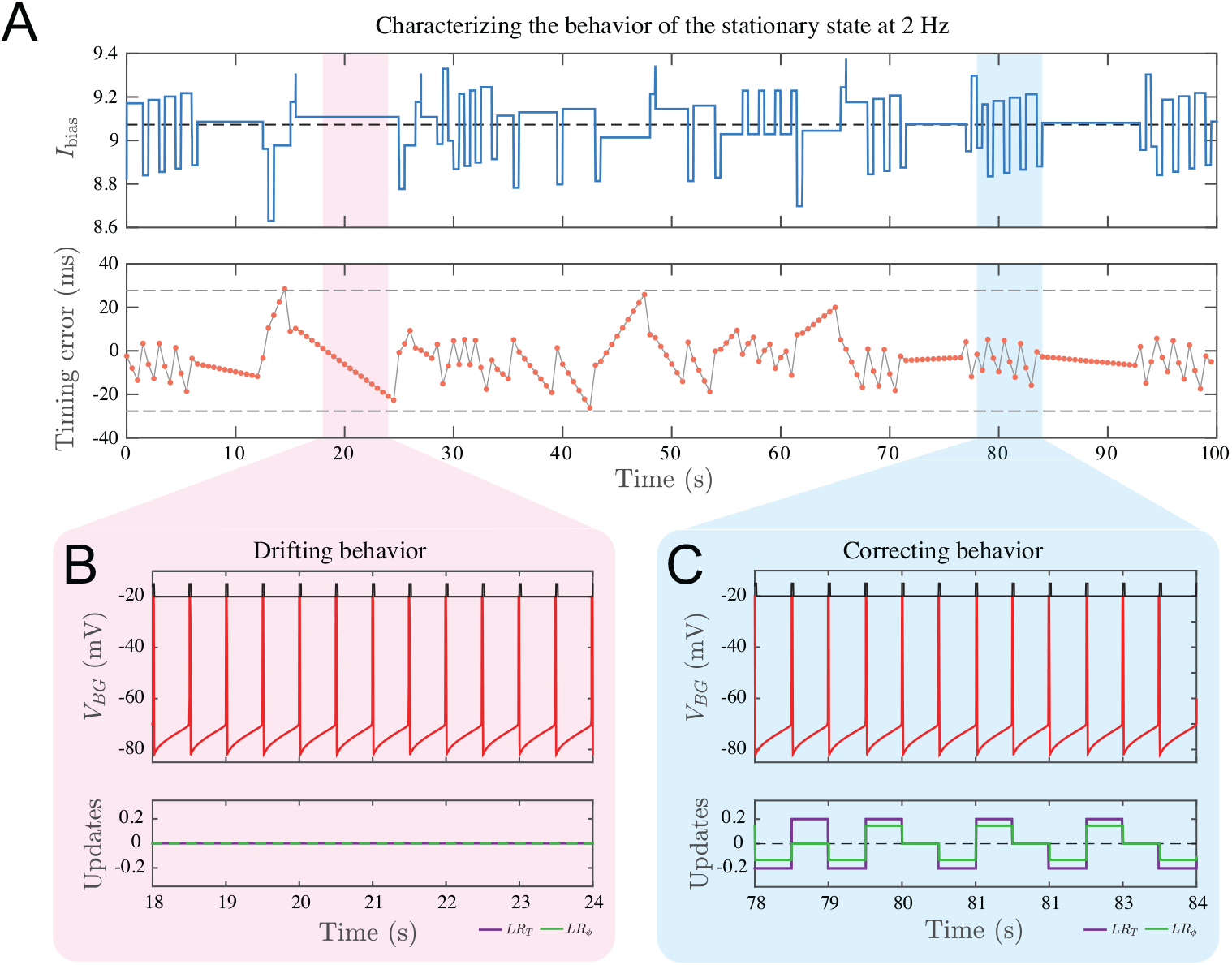
Characteristics of stationary behavior: The *BG* alternates between drifting and corrective behavior while maintaining a beat within one gamma cycle accuracy for a 2 Hz stimulus. **A.** The upper time course of *I*_bias_ shows intervals of rapid change interspersed with intervals of no change. In both cases, *I*_bias_ is centered near *I*_bias_ = 9.06 (dashed line) representing the value that produces exactly 2 Hz oscillations. The lower time course shows the timing error differences, *S* spike time subtracted from *BG* spike times. Intervals of rapid change intermingle with intervals of slower constant change, corresponding to the intervals of rapid changes and no change, respectively, in the upper panel. Timing errors never exceed a time interval of one gamma cycle, ±27.73 ms shown by the dashed grey lines. **B.** Drifting behavior in a 6 s interval shows no updates due to the learning rules during this time. The *BG* spikes systematically advance relative to stimulus onset times, consistent with the negative slope seen in pink in panel A (lower). **C.** Corrective behavior in a different 6 s interval shows how the learning rules *LR_T_* and *LR_ϕ_* help maintain the 2 Hz oscillation. These rules are invoked whenever the counts *γ_BG_* or *CC_BG_* do not match with *γ_S_*.

Although the dynamics of the *BG* are deterministic, they are sensitive in quantitative detail to changes in initial conditions. This is because the learning rules *LR_T_* and *LR_ϕ_* ignore timing differences less than one gamma cycle. To get a more general sense of the fluctuations in *BG* firing times, we ran a simulation for 1000 stimulus cycles and calculated error distribution plots (spike time of *BG* minus spike time of *S*). This was performed at six different stimulus frequencies in steps of 1 Hz (Fig. 6). There are several points to note. First, at all frequencies, the error distribution shows negative mean asynchrony [49,50]. In other words, the actual time of the beat generators firing, on average, preceded the time of the stimulus onset. Second, the variance in the error distribution shows some frequency dependence, particularly with the standard deviation increasing at slower frequencies. Further, the standard deviation increases as the frequency decreases down to 0.5 Hz (simulations not shown). We also found that the standard deviation increases with frequency in the 6-8 Hz range. Accurately tapping at rates above ~ 4 Hz is extremely difficult, hence, no tapping studies exist for this frequency range to either corroborate or contradict our result. However, Drake *et al*. [51] found a U-shaped dependence of the subject’s tempo discrimination ability on frequency in the range 1-10 Hz, consistent with our result. We additionally note that increasing *δ_T_* and *δ_ϕ_* leads to increased variability and larger NMA at all frequencies.

**Fig 6.**
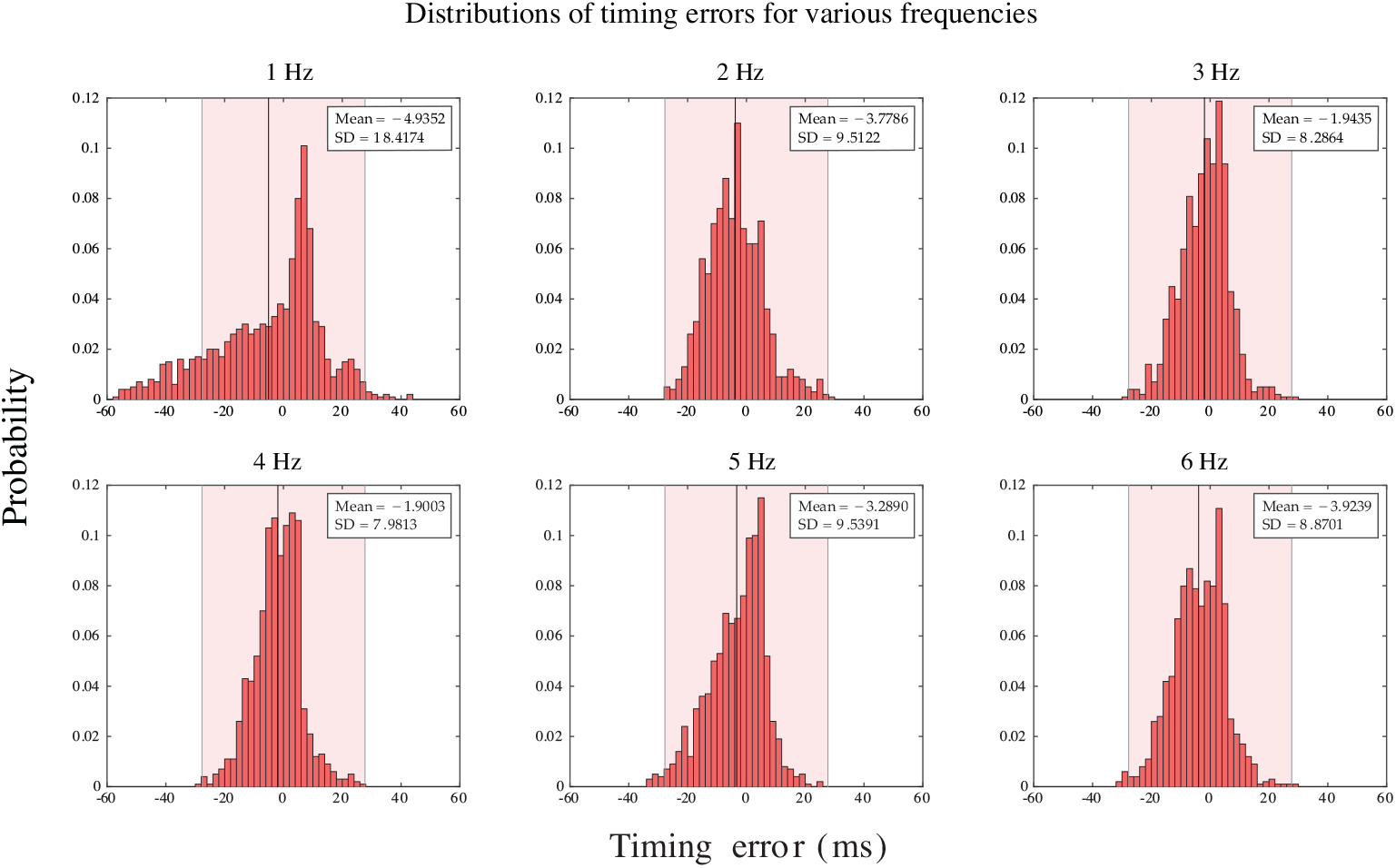
Timing error distributions of beat times. Histograms of timing errors (2 ms bins) based on simulations which ran for 1000 stimulus cycles during stationary behavior at six different stimulus frequencies. Negative mean asynchrony was seen in all cases (black vertical lines) and the standard deviation at the slowest frequency is largest. Pink shading represents a region of one gamma cycle accuracy.

In our model, the interplay of the *BG* neuron’s *f-I* curve and the learning rules, *LR_T_* and *LR_ϕ_*, are responsible for the frequency dependent results. In particular, at both low (~ 0.5Hz) and high (~ 5-8Hz) frequencies the *f-I* curve is steeper than in the intermediate range. Hence, equidistant changes of *I*_bias_ around a mean value result in different changes in instantaneous frequency. The negative mean asynchrony arises from the non-linearity in the *f-I* curve and the asymmetry in the phase learning rule *LR_ϕ_*. As states earlier, *LR_ϕ_* pushes the *BG* to fire before the stimulus.

### Resynchronization time: Responses to frequency changes, phase shifts and temporal deviants

As demonstrated in Fig. 3, the *BG* is able to quickly learn a new frequency. This learning can be quantified as a resynchronization of the *BG*’s spike times with the new stimulus onset times. As previously stated, we declare the *BG* to be resynchronized if three consecutive spikes each fall within one gamma cycle accuracy of an *S* spike. We computed the resynchronization times as a function of several parameters including initial and final stimulus frequency (Fig. 7 shows one example). From a fixed initial stimulus frequency, we changed the stimulus frequency to different values within the range 1 to 6 Hz and computed resynchronization times. In one such case, the stimulus frequency is decreased from 3 to 2 Hz (Fig. 7A). The change is applied at *t* = 0 s (gold star) and the *BG* takes about four seconds (eight stimulus cycles) to synchronize to the new frequency (depicted by the shaded region). During the transient, the learning rules *LR_T_* and *LR_ϕ_* drive *I*_bias_ down in order to slow the *BG* down (lower panels of Fig. 7A). Adjustments due to *LR_T_* occur whenever the *BG* spikes. For the first second after the change in frequency, *BG* spikes at roughly its initial 3 Hz rate. The S neuron spikes at *t* = 0.5 s, which resets *γ_S_*. But the *BG* is not aware of this new larger value of *γ_S_* until it fires at around *t* = 0.66 s. At this point, *γ_S_* > *γ_BG_* and the period learning rule *LR_T_* decreases *I*_bias_. Adjustments due to *LR_ϕ_* occur whenever the stimulus neuron *S* spikes, which now occur at the slower 2 Hz rate. These adjustments depend on the current phase of the *BG* and are seen at times to increase *I*_bias_, but at other times to decrease it. Within two seconds, both rules have succeeded in bringing *I*_bias_ within one gamma cycle accuracy of the 2 Hz target value (dashed black line inside blue band in middle panel). Aligning the spike times then takes a few more seconds. In contrast, an increase in stimulus frequency can lead to much shorter resynchronization times (Fig. 7B). In the transition from a 3 to 4 Hz stimulus frequency, the *BG* only takes about one and a half seconds (six stimulus cycles) to synchronize. The phase learning rule *LR_ϕ_* plays a more prominent role as it is invoked more often due to the increase in stimulus frequency. These examples illustrate two important properties of the resynchronization process. First, the two learning rules act concurrently to adjust I_bias_, but are asynchronous in that *LR_T_* adjustments occur at different times than those of *LR_ϕ_* (see the lower panels of either Fig. 7). Second, adjustments to *I*_bias_ are not periodically applied, they occur at a *BG* or *S* spike, and only the *S* spikes occur periodically.

**Fig 7.**
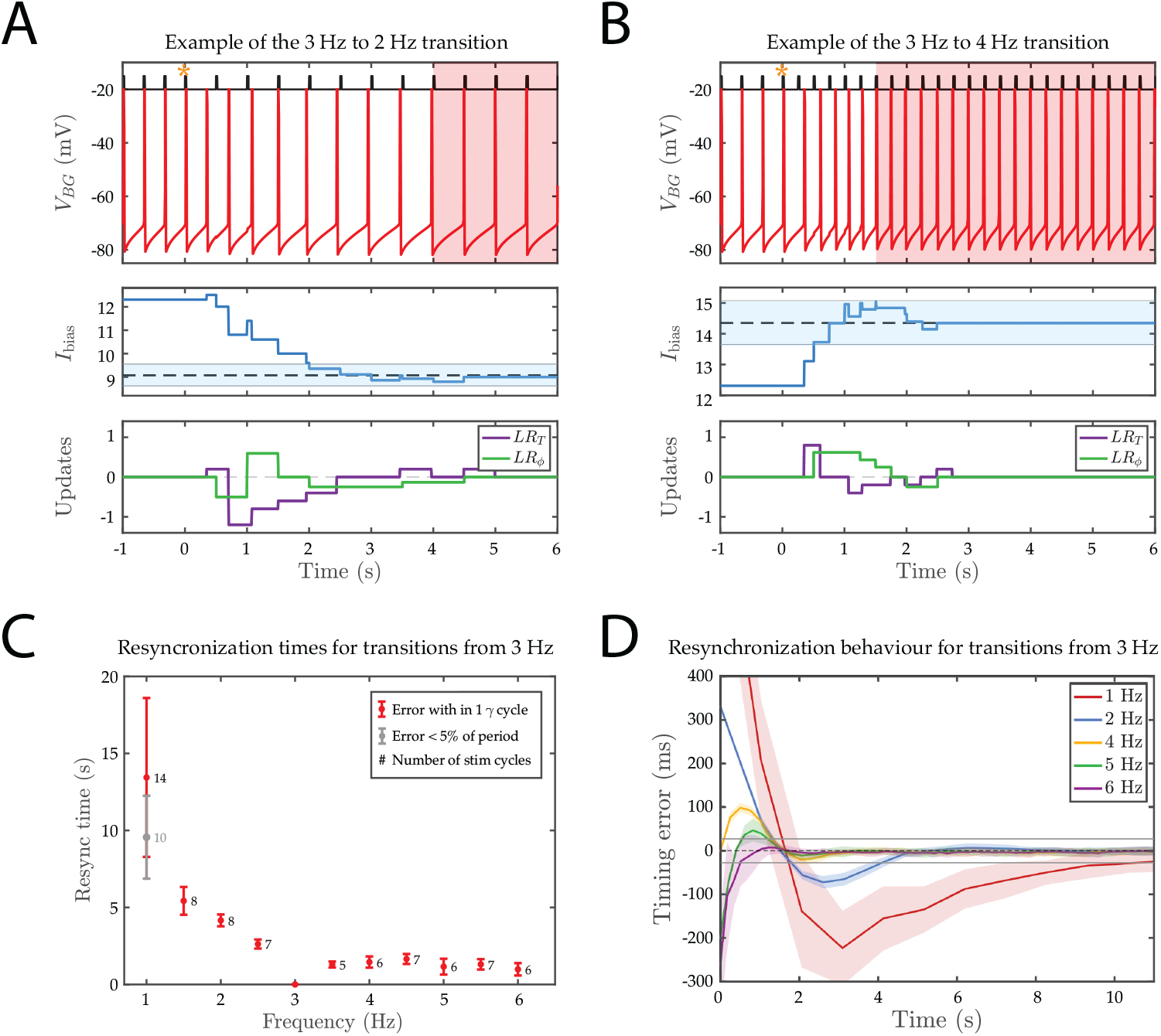
Dynamics of learning a new *IOI_S_*: **A.** and **B.** Top time courses show examples of the resynchronization process of the *BG* spikes with those of *S*. The change in frequency is invoked at *t* = 0 (gold star) and the *BG* resynchronizes within a few seconds (shaded region). Middle and bottom panels show the changes in *I*_bias_ due to *LR_T_* and *LR_ϕ_* indicating that the two rules sometimes change *I*_bias_ in the same direction, but often counteract the effect of the other. *I*_bias_ enters the region of one gamma cycle accuracy(blue shaded band) within a handful of cycles. **C.** Mean and standard deviation for resynchronization times, together with the mean numbers of stimulus cycles needed to achieve the resynchronization are shown. A broader measure of synchrony for the 1 Hz oscillation is also included. The numbers beside the data points indicate the mean resynchronization time in terms of beats. **D.** Mean timing error transition curves for resynchronization to different terminal frequencies averaged over 50 realizations are depicted. For each time course, we aligned the last spike of the *BG* with the last stimulus spike and then subtracted the vector of *S* spike times from the *BG* spike time vector. Shaded bands represent the standard deviation. Grey lines centered about zero depict timing errors within one gamma cycle accuracy. Resynchronization to lower frequencies is longer than to higher ones. Not visible on this scale, the time course for the 1 Hz (red) curve intercepts the y-axis at ~ 850 ms.

Resynchronization times increase with decreasing frequency, but are nearly constant and mostly flat for increasing frequency (Fig. 7C). Decrements from initial to final frequency lead to slower convergence than equally-sized increments. This follows from the slope of the *f-I* curve being steeper while increasing from 3 Hz than when decreasing. For the slowest stimulus frequency (1 Hz) oscillation, we have included a broader measure of synchrony, defined by *BG* spike times falling within 5% of the interonset times, i.e. ± 50 ms around *S* spike times. This definition is consistent with the stationary behavior shown in Fig. 6 where many of the *BG* spikes fall outside of one gamma cycle accuracy. With this broader measure of resynchronization, the average number of cycles and standard deviation of the resynchronization to 1 Hz rhythm are reduced. Although resynchronization times are longer for frequencies decrements, the number of stimulus cycles for resynchronization do not show major differences for increments and decrements, except for the 1 Hz case (the mean number of cycles for resynchronization are reported beside each data point).

The resynchronization process occurs stereotypically depending on whether there is an increase or decrease in frequency (Fig. 7D). We calculated the average cycle-by-cycle time differences for 50 realizations of the resynchronization process from 3 Hz to the target frequency with the standard deviation shown in the shaded region. Decreases (increases) in frequency show initial time errors that are positive (negative). This is due to our spike alignment process (see Fig. 7 caption). Each curve is non-monotonic and, except for the 6 Hz curve, has an under-or over-shoot that transiently takes the curve outside the band of one gamma cycle accuracy (horizontal grey line). The average resynchronization times in Panel C are shown as the time at which the curve reenters this band, i.e. the time at which the timing errors become consistently less than one gamma period. Consistent with our prior results, the standard deviation bands are largest for the 1 Hz curve and relatively similar and small for the other curves.

Resynchronization also occurs when a phase shift of the stimulus sequence occurs. Now consider the 2 Hz case for which the *IOI_S_* is 500 ms. A phase advance will occur if we shorten one *IOI_S_* to be less than 500 ms and then return the remainder of sequence to the original *IOI_S_* of 500 ms. A phase delay is the opposite, where a single *IOI_S_* is elongated. We define the phase *ψ* of the shift to lie within (—0.5,0.5) where negative values represent advances and positive values represent delays. If the phase shift falls within one gamma cycle of the normal onset time, the *BG* is likely to initially ignore it since no change in the gamma counts will occur. But for larger valued phase shifts, resynchronization will need to occur (Fig. 8). As an example, resynchronization for a positive phase shift at *ψ* = 0.4 (Fig. 8A) is much quicker than the corresponding negative phase shift *ψ* = −0.4 (Fig. 8B). The reason for this is how *LR_T_* changes *I*_bias_ in either case. A negative phase shift causes the *BG* to increase its frequency in response to the temporarily shorter *IOI_S_*, followed by a return to a lower frequency. A positive phase shift causes the opposite, a transient decrease in the *BG* frequency followed by an increase. As we have shown earlier, resynchronization times are shorter when the target frequency is larger (Fig. 7). Hence, the model predicts that resynchronization times should be shorter for positive phase shifts (Fig. 8C red). The mean timing errors (standard deviation shaded) for different phase shifts (Fig. 8D) are stereotypical in much the same way as the timing errors for tempo changes. In the current context, the timing errors start out large and then systematically reduce until they fall within one gamma cycle accuracy. The graph clearly shows that the resynchronization after positive phase shifts is faster than after negative shifts, as negative phase shifts exhibit an overshoot.

**Fig 8.**
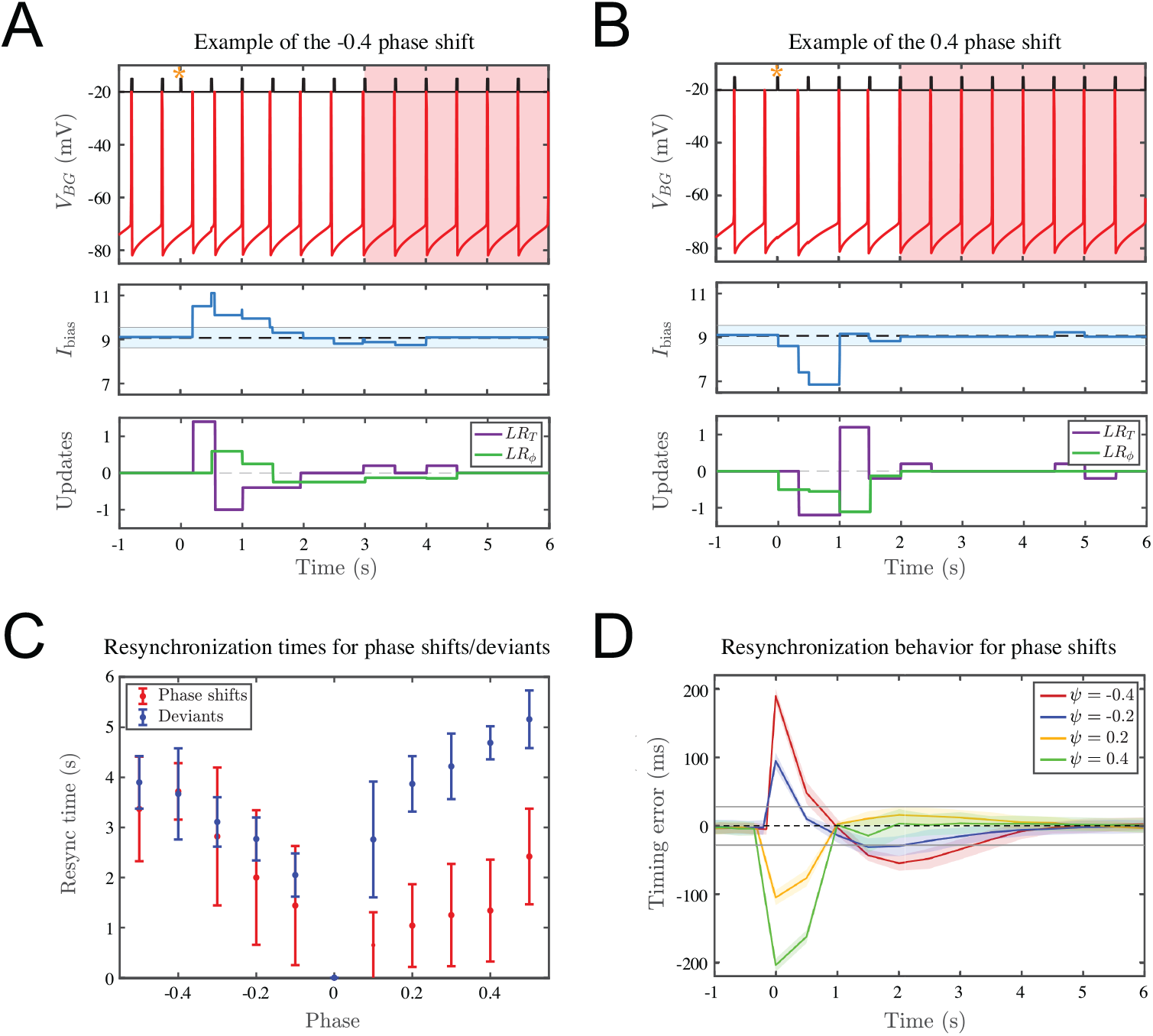
Response to phase shifts and deviants: The description of generic features of each panel is the same as in Fig. 7. **A.** and **B.** Two specific examples show that resynchronization to a positive phase shift is faster than to a negative one. Negative phase shifts cause a transient increase to /bias as shown in the middle panel; positive phase shifts have the opposite effect. After the transient, the return to baseline is faster for the positive phase shift because the *BG* interprets this as equivalent to resynchronization to a higher frequency. **C.** Mean resynchronization times and standard deviations for baseline 2 Hz oscillations are shown for phase shifts (red) and deviants (blue). Notice the asymmetry for both situations, and that resynchronization to a phase shift is generally faster than to a deviant. **D.** Characteristic time courses of average timing differences (standard deviation shaded) for phase shifts give further evidence for the asymmetry in resynchronization times. Note that for the negative phase shifts the model produces an overshoot, where the timing errors temporally fall outside the region of one gamma cycle accuracy (gray line) before returning to it later for eventual resynchronization.

Another case where we see resynchronization is the introduction of a temporal deviant where a single *S* spike occurs at an unexpected early or late time. Unlike phase shifts in which a single *IOI_S_* changes, a single deviant causes both the *IOI_S_* before and the *IOI_S_* after the deviant to change. An early (late) deviant causes a shorter (longer) interonset interval, followed by a longer (shorter) than normal one, followed by a return to the standard *IOI_S_*. The model’s response is different for early versus late deviants. For an early deviant, the phase learning rule *LR_ϕ_* is invoked at the time of the early deviant. This is then followed by the period learning rule *LR_T_* at the next *BG* spike. Both of these signal the *BG* to speed up. For a late deviant, however, the *BG* spikes when it normally would have. At that time, it has no new information about its phase or about the value of *γ_S_*. Therefore there is no change to *I*_bias_. When the late deviant arrives, it now causes *LR_ϕ_* to send a slow down signal to *BG*. But any potential changes due to *LR_T_* have to wait one full *BG* cycle to be invoked. Thus, the model reacts quicker to an early deviant than to a late deviant. Thus we predict shorter resynchronization times for earlier deviants (Fig. 8C blue). Additionally, because of the need to adjust to two different *IOI_S_*s, we predict that resynchronization times due to deviants will be longer than those for comparably sized phase shifts where only a single *IOI_S_* is changed.

### Effects of parameter changes and noise

The model results are robust to perturbations and can operate over a range of parameters. To assess this we ran several simulations (not shown), where we varied intrinsic parameters of the *BG* and the gamma counter speeds. For example, the maximal conductance for *I_L_, I_NaP_* and *I_CaT_* was varied by up to 10 % and we measured the subsequent performance across a range of periods. This did not affect the ability of the *BG* to learn the correct period and phase, because the *f-I* relation remained qualitatively unchanged. At a quantitative level though, the range of Ibias values which yield gamma cycle accuracy will differ and the *BG* neuron may have a different preferred frequency. These results indicate that the *BG* does not require fine tuning of parameters to learn a rhythm.

Next we allowed the speed of the gamma counters for *S* and *BG* to be different. We kept the gamma counter for *IOI_S_* at 36.06 Hz while we varied the gamma counter frequency for the *BG* counter by up to 10%. In all cases, across a range of frequencies there was little qualitative difference in the *BG*’s ability to learn the correct frequency. This is not surprising as the discreteness of the gamma counts allows for similar values of the counts despite there being differences in counting speed. Note that a faster gamma counter for the *BG* tends to lead to earlier firing times relative to stimulus onset times for the parameters we have chosen. In this case, *LR_T_* tends to increase *I*_bias_ since *γ_BG_* is larger than *γ_S_*. On the other hand *LR_ϕ_* decreases the same quantity and it is the parameter-dependent balance between the two rules that determines how much earlier on average the *BG* fires. A slower *BG* gamma counter has the opposite effect. These results imply that the extent of NMA can be manipulated by changing the counter frequency of the *BG* and can even be transformed to a positive mean asynchrony if the *BG* gamma counter is too slow.

To assess the effects of noise, we introduced stochasticity into the gamma counters (see Appendix for details). This acts to jitter the gamma periods, but for modest noise this will only cause the gamma count to discretely change by at most plus/minus 1. Since the *BG* is monitoring its period and phase at each spike and stimulus event, it quickly adjusts to counteract these potential changes. We also see an increase in the standard deviation of the timing error, across all frequencies, during stationary behavior, as well as an increase in NMA. While this widened the distributions (as seen in Fig. 6), approximately 90% of the timing errors remain within one gamma cycle accuracy, apart from at 1 Hz where only 60% of the distribution lies within one gamma cycle accuracy (80% lie within 5% accuracy). Finally, although not explicitly modeled here, one could introduce intrinsic noise in the *BG*, for example a noisy spike threshold or ionic conductance. This small amount of noise would not change the *IBI_BG_* by more that a single gamma cycle and, as above, should not change the *BG*’s ability to synchronize to the external rhythm. In general, noise makes the model fit better with tapping data, exhibiting more variability and larger NMA. Given that human tapping data contains both motor and time keeper noise that our model does not attempt to disentangle (our model does not have an explicit movement component), we did not address this further.

## Discussion

We presented a modeling framework that begins to address how a neuronal system may learn an isochronous rhythm across a range of frequencies relevant to speech and music. We showed how a biophysical conductance-based model neuron, the beat generator (*BG*), adjusts its spiking frequency and aligns its spike times to an external, metronome-like stimulus. Our model employs two gamma frequency oscillators to estimate the number of oscillatory cycles between certain salient events. We posit a mechanism that compares the states of these independent counts to inform the *BG* to either increase or decrease its instantaneous frequency and adjust its relative phase. With this idealized paradigm, we showed that the *BG* quickly learns to hold a beat over a range of frequencies that includes, but is not limited to, 1 to 6 Hz. Further, we showed how the *BG* reacts within a few cycles to changes in tempo, phase shifts (permanent realignment of the stimulus sequence) and the introduction of deviants (temporary misalignment of a single stimulus event). Of particular note, the *BG* displays an asymmetry in reacting to changes to the rhythm. It adapts more quickly when the tempo is increased as opposed to decreased; correspondingly, it reacts faster to phase delays than phase advances, but slower to late deviants than early deviants. Importantly in our model formulation no direct input from the stimulus to the *BG* is provided. This implies that the *BG* is *learning* the correct period and phase rather than being *entrained* to them. Secondly, no explicit or exact time intervals are required to be calculated, implying that the *BG* does not need specific mechanisms to exactly track time. Instead, in order to tune the *BG*, one needs only to know, in some rough sense, whether the *BG*’s spikes are happening too fast or slow relative to stimulus frequency and too early or late relative to the stimulus onset. Finally, because of the discrete nature of the gamma counters the *BG* dynamics are robust to modest parameter changes and noise.

### Beat generation differs from beat perception

Beat perception as described in many previous studies [52,53] refers to the ability of an individual to discern and identify a basic periodic structure within a piece of music. Beat perception involves listening to an external sound source as a precursor to trying to discern and synchronize with the beat. Alternatively, we might ask how do we (humans) learn and then later reproduce a beat in the absence of any external cues. Such issues and questions lead us to consider what neuronal mechanisms might be responsible for producing an internal representation of the beat. At its most basic level, we refer to this as beat generation, and a neuronal system that does so we call a beat generator. Different than beat perception, beat generation is envisioned to be able to occur in the absence of an external cue. A *BG* is a neural realization of an internal clock that can be used as a metronomic standard by other internally driven processes that depend on time measurements. While demonstration of a beat involves a motor action (tapping, clapping, vocalizing, head bobbing), the *BG* could include a general representation of a motor rhythmicity but the specific motor expression (say, foot tapping) may not be an integral part of the *BG*.

### Learning a beat differs from entraining to a beat

Our formulation proposes that time measurement for beat perception and the beat generator model are oscillator-based. In this view a beat can be learned and stored as a neuronal oscillator (cell or circuit). The frequency range of interest, 1-6 Hz, is relatively low compared to many other neuronal rhythms, but similar to those seen in sleep. We rely on faster (gamma-like) oscillators to provide clocklike ticks and we assume two counters and a comparator circuit can be used for adjusting the *BG* period and phase to match with the stimulus. Conceptually, counting and comparing with a target period are essential features of the algorithmic (or sometimes called, information processing timekeeper) approach falling into the class of error-correction strategies; see [16, 17, 20–22, 54] for examples of two-process models. These models suggest mechanisms used by humans to bring their movements into alignment with a rhythmic stimulus. They do not, however, provide a biological framework for these mechanisms. We provided a neuronal implementation of the *BG* in the form of an oscillator with a tunable biophysical knob and two learning rules; the *BG* is a continuous-time dynamical system, a realizable neuronal oscillator. It does not require a separate reset mechanism. The implementation also does not require a separate knob for phase correction; the two learning rules both make adjustments/corrections to the same parameter, *I*_bias_, and they are ongoing whenever a stimulus is present. We propose this *BG* as the internal clock — an oscillator that learns a beat and keeps it.

A different class of oscillator models for beat perception relies on large networks of neuronal units [12, 24, 28]. The units’ intrinsic frequencies span the range of those that are relevant in speech and music. In the neural entrainment models of Large and collaborators, different units within the ensemble respond by phase-locking to the periodic stimulus. Units with intrinsic frequencies near that of the metronome will entrain 1:1 while those with higher intrinsic frequencies entrain with different patterns, such as 2:1. Dominant responses are found at harmonics and sub-harmonics of the external input. Amplitude, but not precise timing relative to stimulus features (say, stimulus onset times), are described in these models. The framework is general although the identities of neuronal mechanisms (synaptic coupling or spike generation) are not apparent as the description is local, based on small amplitude perturbation schemes around a steady state and the coupling is assumed to be weak. The approach is nonlinear and provides interpretations beyond those of linear models, e.g. it identifies a beat for complex input patterns even if the beat/pulse is not explicitly a component of the stimulus [12].

Our model cannot be described as entrainment in the classical sense. Entrainment occurs when an intrinsically oscillatory system is periodically forced by an external stimulus to oscillate and, in the present context, to phase lock at the forcing frequency (or some subharmonic) that may differ from its endogenous frequency. Our *BG* neuron is not entrained by the stimulus but rather it learns the frequency of the stimulus. The *BG*’s frequency is adapted indirectly through the control parameter in order to match with the stimulus. The influence of the stimulus on the *BG* diminishes as learning proceeds. In fact, in the continuous time version when the frequency and phase are eventually learned, the *BG* no longer requires the stimulus; it will oscillate autonomously at the learned frequency if the stimulus is removed or until the stimulus properties change. In the discrete time version, even after the stimulus and *BG* periods and phase agree (to within a gamma period accuracy) modest adjustments are ongoing to maintain the rhythm. In contrast, for an entrainment model, the oscillator’s parameters are fixed. The stimulus does not lead to a change in the oscillator’s intrinsic properties. For a transient perturbation, the dynamics of resynchronization are according to an entrainment unit’s phase response curve, which instantaneously changes the current phase of the oscillator. In contrast, the *BG*’s response to transient inputs impacts the parameter *I*_bias_ invoked by adjustments according to either or both of the period and phase learning rules. Our model is further distinguished from entrainment models in that the *BG* strives for zero phase difference but in an entrainment setting there is typically a phase difference between the stimulus and the units. Finally, for an entrainment model the coupling from stimulus to oscillator is periodic. In our model the influence of a periodic stimulus is delivered both periodically (via *LR_ϕ_*) and aperiodically (via *LR_T_*).

### Implications of using gamma counters

Although humans can learn to accurately estimate time intervals [1], little is known about the neural mechanisms used to generate these estimates. For beat generation we are positing an ability to estimate time intervals (e.g., between stimulus onset events) in real time in an ongoing and flexible manner. We introduced the idea of gamma counters to perform such measurements. These counters provide a rough estimate of elapsed time that can be used to compare the internal representation of an interval with that of an external cue. The model then produces a finer representation of the interval by adjusting the *BG*’s spike time and period. There is growing evidence for the existence of counting mechanisms within neuronal systems. For example, Rose and collaborators have demonstrated that neurons in the auditory mid-brain of anurans (frogs and toads) count sound pulses in order to make mating decisions [36, 55]. These neurons have been called ‘interval counting neurons’ because they respond only after a threshold number of pulses have been counted provide that those pulses are spaced in time intervals of specific lengths [37]. In a very different context, it has been recently demonstrated that mossy fiber terminals in rat hippocampus have the ability to count action potentials, an ability cited as improving the reliabilty and accuracy of information transfer [38].

The discreteness of the gamma counter, used in our model, leads to variability in the *BG* spike times, allowing the model to exhibit negative mean asynchrony (NMA). This is consistent with finger tapping experiments which show that humans display variability in their tap times relative to an isochronous stimulus and tend to, on average, tap before the stimulus [13]. As discussed earlier, the NMA as shown in Fig. 6 is rather modest, however, changes in parameters will lead to larger NMA. In contrast, replacing the gamma frequency oscillators with continuous time clocks, which exactly determine time intervals, leads to perfect phase alignment, *ϕ* = 0 (no NMA). Thus, our work posits the existence of discrete time clocks as a potential source of intertap time variability.

The gamma counters also provide an upper bound on the stimulus frequency which can be reliably learned by the *BG* neuron. For the 36 Hz clocks used here, this limit is roughly 9 Hz. After this point the phase rule overcorrects, transiently increasing *I*_bias_ to a value corresponding to a much larger frequency. We stress that this upper bound is dependent on the specific gamma frequency, and faster clocks may be used to keep track of shorter intervals. An interesting experimental study would be to look at the EEG power spectrum while subjects listen to periodic stimuli and monitor whether the gamma band activity changes with stimulus presentation rate.

### Relation to interval timing and other models for beat production

Many interval timing models involve accumulation (continuous time or counting of pacemaker cycles) with adjustment of threshold or ramp speed [6, 7] to match the desired time interval. Applications to periodic beat phenomena, say the metronome case, would include instantaneous resetting and some form of phase adjustment/correction [56,57]. Algorithmic models may not specifically identify the accumulator as such, but instead refer to counters or elapsed time. Our *BG* model shares some features with interval models for beat production (as described in [9] and [58]), as the *BG* relies on counters and accumulators. Additionally, as described earlier, it shares features with entrainment models, as the *BG* is a nonlinear oscillator. In short, the *BG* is a hybrid.

Interval- and oscillator-based models are related. Even if not explicitly stated as such, in an interval model, the accumulator and its reset are equivalent to highly idealized models for neuronal integration, the so-called integrate-and-fire (*IF*) class of models [59]. For steady input, the state variable rises toward a target value (that is above the event threshold), rising linearly for a non-leaky *IF* model and with a decreasing slope for a leaky *IF* model (*LIF*), and is reset once the state variable exceeds the threshold. These *IF*/*LIF* models are dynamical system oscillators, and are also nonlinear by way of the reset mechanism. However, the time constant/integration rate required for beat applications is much longer/slower than in typical applications of *IF* models for neuronal computations where timescales of 10-30 ms are more common. These models have entrainment and phase-locking properties [60, 61] and they typically show a phase difference from the stimulus. Extended in this way, periodic in time, such an interval model can be recast as an entrainment model (see also [39]). As noted by Loehr *et al*. [39], differences between such interval and continuous oscillator models do appear in some circumstances. Adding a plasticity mechanism, say for the threshold or input drive, then allows learning of a period. We described how one may analyze the dynamics of such an *LIF* oscillator-like interval model in terms of a map (Fig 4). One could additionally add a phase correction mechanism as in two-process models in order to achieve zero-phase difference. This can be achieved in a *LIF* model, for example, by adjusting the reset condition after reaching threshold or by utilizing phase response curves. Our mechanism for phase correction differs from these approaches in that we target the excitability parameter *I*_bias_ for adjustment. This has the advantage that the *BG* learns the correct phase and period allowing it to continue to hold a beat after the stimulus is removed, similar to other two-process interval models.

The effects of noise on time estimation/production have been studied with interval models, cast as first passage time problems for accumulator models (drift-diffusion models) [62–64]. In that context, the issue of scalar timing is of significance [5, 63, 65, 66], however the time intervals of interest are typically longer than what one would find in a musical context. Wing and Kristofferson [16,17] considered effects of noise and contrasted sensory noise with motor sources of noise, concluding that timekeeper noise was frequency dependent but motor noise was not. Whether or not scalar timing holds for short rhythmic intervals is unsettled. A number of tempo discrimination studies have failed to produce any evidence for frequency dependent errors for periods below 1000 ms [51,67]. However, Collyer *et al*. [68] reports scalar timing in the distribution of tap times when tapping to an isochronous rhythm.

A distinctive interval model was developed by Matell and Meck [69,70] – the striatal beat frequency (SBF) model. In this neuromechanistic description, the basic units are neuronal oscillators with different fixed frequencies. All oscillators are reset at *t* = 0; differences in frequencies of convergent units will eventually lead to collective near-coincidence (so-called beating phenomenon of non-identical oscillators) at a time that through learned choices (synapses onto coincidence detector units) can match the desired interval. It may be extended to the periodic case and considered for beat generation as discussed in [56,57] although the brain regions involved may be different for explicit time estimation than for rhythmic prediction/reproduction [71,72].

### Limitations of this *BG* model

We consider here only the case of isochronous inputs. A natural next step is to consider more complex, non-isochronous stimulus sequences. Additionally, we have side-stepped questions of perception in order to focus solely on timing. Our *BG* model does not recognize variations in pitch or sound level. For example, if stimulus events were alternating in, say, sound level, (as in [73]) our model, as is, would not capture the effects. An extension of our model involving pairs of stimulus and beat generator clocks for each sound level could conceivably address this shortcoming.

We have chosen a particular biophysical instantiation for the *BG*. The capabilities of learning and holding a beat over a range of frequencies depends only on the monotonic frequency dependence of the control (“learnable”) parameter and would not be compromised by variation of biophysical parameters. Some features of the *BG* dynamics (say, the degree and signatures of asymmetries in resynchronization for speeding up or slowing down) can be expected to depend on the specifics of, say, the relationship between *I*_bias_ and the intrinsic frequency, but we have not explored this in detail.

The learning rules *LR_T_* and *LR_ϕ_* utilized in our study both target the excitability parameter *I*_bias_ with a simple goal to either speed up or slow down the *BG* so that it synchronizes with the stimulus. Alternatively, the drive could be provided as the summed synaptic input from a population of neurons afferent to the *BG*. The synaptic weights onto the *BG* and/or internal to the afferent population could be plastic and affected by our learning rules which in spirit are similar to spike time dependent plasticity rules [74]. Our model assumes significant increments of drive at each learning step, leading to fast learning. This may be relatable at a population scale to balanced network models, where fast learning can be achieved with smaller step changes due to the large number of synapses [75]. Currently, during synchronization continuation, our *BG* model retains its estimate of the most recent stimulus period, *γ_S_*. We have not yet included a slow decay of this memory or a slow degradation of the *BG* rhythm. It is plausible that the addition of noise could lead to this slow drift after the stimulus is removed since, as we showed, noise does introduce additional variablity during stationary behavior.

We have not ascribed a location for the *BG* within a specific brain region. As a result, we have not addressed issues of sensorimotor synchronization (SMS) where sensory processing of a beat must be coordinated with the motor action that demonstrates the beat (e.g. finger tapping). Several models for SMS in the context of beat perception already exist, for example the two-layer error-correction model of Vorberg and Wing [76] and the entrainment model of Large *et al*. [12] described earlier. Van der Steen and Keller have developed the Adaptation and Anticipation Model (ADAM) [22], a type of algorithmic error-correction SMS model, and they noted a need for an extended ADAM that would incorporate dynamical systems principles. Our model could certainly be a starting point for such an endeavor. Patel and Iversen [77] proposed the Action Simulation for Auditory Prediction (ASAP) hypothesis. In their conceptual model, the motor system primes the auditory system to be able to process auditory input. In particular, ASAP proposes that the motor system is required for beat perception. Generally, these studies raise questions about whether the causal roles of sensory and motor systems can be disambiguated in the context of beat perception and beat generation. Addressing such questions from a modeling perspective is a natural next step.

### Predictions based on the *BG* model

Our model framework allows us to make several predictions, which are summarized here. First, the *BG* model succeeds at synchronization continuation [78]; it can hold a beat after the sound stimulus terminates. The *BG* will continue to oscillate with fine adjustments of its period as needed, according to *LR_T_*, to match that of the most recently stored *IOI_S_* of the stimulus (as in Fig. 3). Error-correction models would also continue oscillating at the last stored frequency [20, 21]. In contrast, for an entrainment model without a plasticity mechanism, the oscillators are likely to return to their original intrinsic frequencies after the stimulus is removed. However, Large *et al*. [12] have illustrated using their two-layer model, that the network can hold a beat if the units within the motor layer, have bistable properties, i.e. a unit may have a steady state (damped resonator) coexistent with an oscillator state. The main difference between our approach and others is that period error correction (*LR_T_*) occurs at the *BG* spike times rather than at stimulus onsets. As such, even after stimulus removal, comparison between the *BG* period and last stored stimulus period continues. Second, the time course of adjusting to a sudden tempo change occurs over seconds and has more or less monotonic phases of slowing down or speeding up (Fig. 7). If the new sound stimulus is stopped during this transition, we predict from the model that the *BG* will still learn the new beat frequency. However, the phase of the *BG* will differ depending on when during the transition the stimulus is removed. This could be detected using EEG or perhaps a finger-tapping demonstration. This prediction differs from those made by traditional error-correction models which will cease making updates after the stimulus is turned off. Hence, these models may not reach the new frequency, and instead settle at an intermediate frequency. An entrainment model also relies on the external input to match its frequency. Thus if the sound stimulus is stopped during the transition, an entrainment model is likely to return to its basal frequency, and not learn the new one. Third, resynchronization should be faster after a phase shift of the rhythmic stimulus than after a single timing-deviant sound event. Lastly, the model predicts an asymmetry in the resynchronization process after phase shifts (advance versus delay), deviants (early versus late) and tempo changes in the stimulus sequence. We hesitate to assign whether the resynchronization time will be shorter or longer in these cases, as this property depends on the intrinsic dynamics of the *BG* as defined by its *f*-*I*_bias_ relationship. Similarly, the specific responses of an entrainment model for these cases depends on the phase response properties of the underlying oscillator model. Typically these are one time adjustments to the phase of the oscillator, followed by a transient return to the entrained solution. Thus rather than elucidating concrete differences in predictions of our *BG* model versus others, instead consideration of different models may help to narrow the choice of biophysical currents that produce *I*_bias_-*f* or phase response properties that allow neuron spike times to match empirical data.

### Future directions

There are several questions that we plan to address in our future modeling and behavioral studies. How sensitively do timing errors depend on variability of the gamma counters and, say, on stimulus frequency? To what degree can the *BG* model track modulations of the beat frequency? Different candidate beat generators that possess different ionic currents will have *f-I* relationships that are different than the ones presented here. How sensitively do the quantitative results described here depend on the shape of these curves? When considering an ensemble of beat generating neurons, how does coupling between these neurons shape the dynamics of learning? How could the model be enhanced to become predictive, to not just track modulation but to predict dynamic trends? Going beyond isochronous timing only, we plan to consider more complex rhythms. For example, suppose we consider the effect of shifting identically the timing of alternate stimulus tones. Eventually, after a sequence of modest shifts, the beat frequency would be halved although the number of stimulus events would be maintained but with a different temporal pattern. How is the transition of frequency halving executed dynamically? Perhaps there is a regime of shift values where beat determination is ambiguous, a possible regime of bistability. A different manipulation toward a complex stimulus could involve parametrically changing the sound intensity or pitch of alternate tones. Such cases will bring us toward questions of perception and auditory streaming together with beat perception.

### Conclusion

The questions surrounding how we perceive and keep a beat are easy to pose but developing models for beat perception and generation present challenges. Our model is a first-pass attempt at formulating and analyzing a neuromechanistic model that can learn a beat. Our approach stems from a neurobiological and dynamical systems perspective to develop neuronal system-based models for beat learning and generation. The essential features involve neuro-based elapsed timekeepers, time difference comparators and a neural oscillator (cellular or circuit level) with some plasticity and learning rules. Looking ahead one hopes for development of more general beat and rhythm pattern generators (for complex rhythmic sounds, music pieces) that can be stored in a silent mode and are both recallable and replayable.

## Supporting information

### S1 Appendix

The model for the *BG* consists of a set of biophysical conductance-based equations which we call the *I_NaP_* model that incorporate a low-threshold calcium current *I_CaT_*, a sag current, *I_h_*, a persistent sodium current *I_NaP_* and a leak current, *I_L_*. The current balance equations are given by

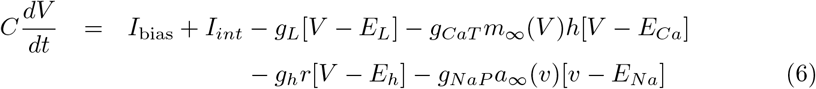

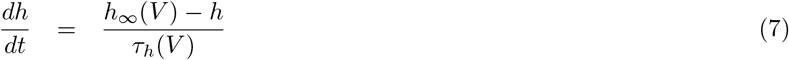

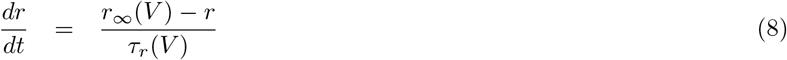

The term *I*_bias_ refers to a drive whose value determines whether the isolated *BG* can oscillate and if so, at which frequency. We set *I_int_* = − 33*μ*A/cm^2^ so that if *I_bias_* = 0*μ*A/cm^2^ there are no oscillations. The parameters *g_CaT_, g_h_, g_NaP_, g_L_* and *E_Ca_, E_h_, E_Na_, E_L_* refer to conductances and reversal potentials for the calcium, sag, sodium and leak currents, respectively. The functions *x*_∞_(*V*) = 1/(1 + exp(−(*V* − *v_x_*)/*k_x_*)) for *x* = *m, a, r, h* are each sigmoidal functions with half-activation voltages *v_x_* and accompanying slopes *k_x_*. The time constants are *τ_h_*(*V*) = *τ_L_*/(1 + exp((*V* − *v_h_*)/*k_h_*)) + *τ_R_*(1 + exp(−(*V* − *v_h_*)/*k_h_*)) and *τ_r_*(*V*) = *τ_r_max__*/ cosh((*V* − *v_r_τ__*)/(2*k_r_τ__*)). The T-current is considered to have instantaneous activation, modeled by *m*_∞_(*V*) and a slow inactivation, governed by the *h* variable. The sag current simply has a slow activation variable *r*. The persistent sodium current has just instantaneous activation given by *a*_∞_(*V*). The parameter values are as follows: *C* =1, *g_CaT_* = 11, *g_h_* = 1, *g_NaP_* = 0.1, *g_L_* = 1.6, *E_Ca_* = 50, *E_h_* = −30, *E_Na_* = 50, *E_L_* = −70, *v_m_* = −40, *k_m_* = 6.5, *V_a_* = −67, *k_a_* = 1, *v_r_* = −70, *k_r_* = 12, *v_h_* = −60, *k_h_* = 6, *v_r_τ__* = −75, *k_r_τ__* =8, *τ_L_* = 30, *τ_R_* = 5, *τ_r_max__* = 850 (where capacitance is in units of *μ*F/cm^2^, conductances are in mS/cm^2^, time constants are in ms and all others parameter units are mV):.

The equations for the *S* neuron are

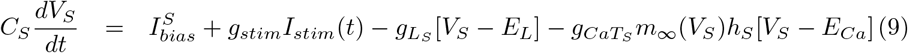

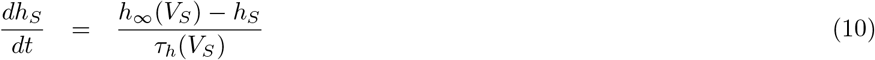

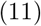

The term *I_stim_*(*t*) is the periodic current provided from the stimulus. During each cycle, it is positive for 25 ms and 0 otherwise. It is taken to be large enough to ensure that *S* fires within 5 ms of sound onset times. The *m*_∞_, *h*_∞_ and *τ_h_* functions are as above, the parameter values are the same unless otherwise stated: 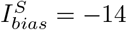, *g_stim_* = 6, 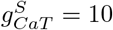 (units as above).

The *γ* counters are constructed as follows. Solve *x*′ = −*x*/*τ_x_* with *x*(0) = 2 until it reaches 1 at t = *t_g_* and is reset to 2; 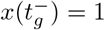 reset to 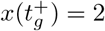. The counters keep track of the number of resets. We chose *τ_x_* = 40 ms which yields an inter-spike interval of 27.73 ms (frequency of 36.06 Hz). Heterogeneity between the *BG* and *S* counters of roughly 10% was introduced to the *IOI_S_* by varying *τ_x_*. In the case of stochasticity, Ornstein-Uhlenbeck noise was added to the *x* variable, with a timescale of 5 ms and Gaussian white noise with mean 0 and standard deviation 0.005. All numerical simulations were carried out in MATLAB.

## Acknowledgments

A. Bose thanks the Courant Institute of Mathematical Sciences at New York University for their support during his sabbatical. A. Byrne was supported by the Swartz Foundation.

